# An Evolutionarily Conserved piRNA-producing Locus Required for Male Mouse Fertility

**DOI:** 10.1101/386201

**Authors:** Pei-Hsuan Wu, Yu Fu, Katharine Cecchini, Deniz M. Özata, Amena Arif, Tianxiong Yu, Cansu Colpan, Ildar Gainetdinov, Zhiping Weng, Phillip D. Zamore

**Affiliations:** Howard Hughes Medical Institute and RNA Therapeutics Institute, University of Massachusetts Medical School, 368 Plantation Street, Worcester, MA 01605, USA; Bioinformatics Program, Boston University, 44 Cummington Mall, Boston, MA 02215, USA; Program in Bioinformatics and Integrative Biology, University of Massachusetts Medical School, 368 Plantation Street, Worcester, MA 01605, USA; Department of Biochemistry and Molecular Pharmacology, University of Massachusetts Medical School, 365 Plantation Street, Worcester, MA 01605, USA; School of Life Sciences and Technology, Tongji University, Shanghai, China

**Keywords:** PIWI-interacting RNA, piRNA, MIWI, A-MYB, MYBL1, spermatogenesis, acrosome, zona pellucida, sperm, pachytene piRNA, meiosis

## Abstract

Pachytene piRNAs, which comprise >80% of small RNAs in the adult mouse testis, have been proposed to bind and regulate target RNAs like miRNAs, cleave targets like siRNAs, or lack biological function altogether. Although piRNA pathway protein mutants are male sterile, no biological function has been identified for any mammalian piRNA-producing locus. Here, we report that males lacking piRNAs from a conserved mouse pachytene piRNA locus on chromosome 6 (*pi6*) produce sperm with defects in capacitation and egg fertilization. Moreover, heterozygous embryos sired by *pi6*^−/−^ fathers show reduced viability in utero. Molecular analyses suggest that *pi6* piRNAs repress gene expression by cleaving mRNAs encoding proteins required for sperm function. *pi6* also participates in a network of piRNA-piRNA precursor interactions that initiate piRNA production from a second piRNA locus on chromosome 10 as well as *pi6* itself. Our data establish a direct role for pachytene piRNAs in spermiogenesis and embryo viability.

**Highlights:**

- Normal male mouse fertility and spermiogenesis require piRNAs from the *pi6* locus
- Sperm capacitation and binding to the zona pellucida of the egg require *pi6* piRNAs
- Heterozygous embryos sired by *pi6*^−/−^ fathers show reduced viability in utero
- Defects in *pi6* mutant sperm reflect changes in the abundance of specific mRNAs.

## INTRODUCTION

Only animals produce PIWI-interacting RNAs (piRNAs), 21–35-nt RNAs that form the most abundant small RNA class in the germline. piRNAs protect the germline genome from transposons and repetitive sequences, and, in some arthropods, piRNAs fight viruses and transposons in somatic tissues (Houwing et al., 2007; Aravin et al., 2008; Batista et al., 2008; Das et al., 2008; Lewis et al., 2018). The mammalian male germline makes three classes of piRNAs: (1) 26–28 nt transposon-silencing piRNAs predominate in the fetal testis (Aravin et al., 2008); (2) shortly after birth, 26–27 nt piRNAs derived from mRNA 3′ untranslated regions (UTRs) emerge (Robine et al., 2009); and (3) at the pachytene stage of meiosis, ∼30 nt, non-repetitive pachytene piRNAs appear. Pachytene piRNAs accumulate to comprise >80% of all small RNAs in the adult mouse testis, and they continue to be made throughout the male mouse reproductive lifespan. These piRNAs contain fewer transposon sequences than the genome as a whole, and most pachytene piRNAs map only to the loci from which they are produced. The diversity of pachytene piRNAs is unparalleled in development, with >1 million distinct species routinely detected in spermatocytes or spermatids. Intriguingly, the sequences of pachytene piRNAs are not themselves conserved, but piRNA-producing loci have been maintained at the syntenic regions across eutherian mammals (Girard et al., 2006; Chirn et al., 2015), suggesting that the vast sequence diversity of pachytene piRNAs is itself biologically meaningful.

In mice, 100 pachytene piRNA-producing loci have been annotated (Girard et al., 2006; Grivna et al., 2006; Lau et al., 2006; Ro et al., 2007; Li et al., 2013). All are coordinately regulated by the transcription factor A-MYB (MYBL1), which also promotes expression of proteins that convert piRNA precursor transcripts into mature piRNAs, as well as proteins required for cell cycle progression and meiosis (Bolcun-Filas et al., 2011). Of the 100 piRNA-producing loci, 15 pairs of pachytene piRNA-producing genes are divergently transcribed from bidirectional, A-MYB-binding promoters (Li et al., 2013). The contribution of pachytene piRNAs from each piRNA-producing locus is unequal, with just five loci located on five different chromosomes—*pi2*, *pi6*, *pi7*, *pi9*, and *pi17*— contributing >50% of all pachytene piRNA production at 17 days postpartum (dpp).

Loss of proteins required to make pachytene piRNAs, including the pachytene piRNA-binding protein, MIWI (PIWIL1), invariably arrests spermatogenesis without producing sperm, rendering males sterile (Deng and Lin, 2002; Reuter et al., 2011; Zheng and Wang, 2012; Li et al., 2013; Castañeda et al., 2014; Wasik et al., 2015). Yet loss of the chromosome 17 pachytene piRNA-producing locus, *17-qA3.3-27363(−),26735(+)* (henceforth, *pi17*), has no detectable phenotype or impact on male fertility (Homolka et al., 2015), even though *pi17* produces ∼16% of all pachytene piRNAs in pachytene spermatocytes. Similarly, mice with disrupted expression of a chromosome 2 piRNA locus are viable and fertile (P.-H.W., K.C., and P.D.Z, unpublished; Xu et al., 2008). Consequently, the function of pachytene piRNAs in mice is actively debated. One model proposes that pachytene piRNAs regulate meiotic progression of spermatocytes by cleaving mRNAs during meiosis (Goh et al., 2015; Zhang et al., 2015). Another posits that pachytene piRNAs direct degradation of specific mRNAs via a miRNA-like mechanism involving mRNA deadenylation (Gou et al., 2014). A third model proposes that MIWI functions without piRNAs, and that piRNAs are byproducts without a critical function (Vourekas et al., 2012). Compelling evidence supports each model.

In fact, direct demonstration of piRNA function in any animal has proven elusive. Only two piRNA-producing loci have been directly shown to have a biological function; both were identified genetically before the discovery of piRNAs and are found only in members of the melanogaster subgroup of flies (Livak, 1984; Livak, 1990; Palumbo et al., 1994; Pélisson et al., 1994; Bozzetti et al., 1995; Prud’homme et al., 1995; Robert et al., 2001; Robert et al., 2001; Mével-Ninio et al., 2007; Chirn et al., 2015). In male flies, piRNAs from *Suppressor of Stellate*, a multi-copy gene on the Y chromosome, silence the selfish gene *Stellate*; deletion of *Suppressor of Stellate* leads to Stellate protein crystals in spermatocytes (Aravin et al., 2001; Aravin et al., 2003). In female flies, deletion of the piRNA-producing *flamenco* gene, which is expressed in somatic follicle cells that support oogenesis, leads to *gypsy* family transposon activation and female infertility (Brennecke et al., 2007; Saito et al., 2009).

Here, we report that a promoter deletion in the chromosome 6 pachytene piRNA locus *6-qF3-28913(−),8009(+)* (*chr6: 127,776,075–127,841,890*, mm10; henceforth, *pi6*) that eliminates *pi6* piRNA production disrupts male fertility. The *pi6* locus, one of the five most productive piRNA-producing loci in mice, generates 5.8% of pachytene piRNAs in the adult testis and is conserved among eutherian mammals. Mice lacking *pi6*-derived piRNAs produce normal numbers of sperm and continue to repress transposons. However, *pi6* mutant sperm fertilize eggs poorly due to defective sperm capacitation and fail to penetrate the zona pellucida, a glycoprotein layer surrounding the egg. Consistent with these phenotypes, the steady-state abundance of mRNAs encoding proteins crucial for sperm acrosome function and penetration of the oocyte zona pellucida is increased in spermatids from *pi6* males. In addition to decreasing specific mRNA abundance, *pi6* piRNAs concurrently facilitate biogenesis of piRNAs from other loci. Our findings provide direct evidence for a biological function for pachytene piRNAs in male mouse fertility, and *pi6* promoter deletions provide a new model for future studies of piRNA biogenesis and function.

## RESULTS

### *pi6* Promoter Deletion Eliminates *pi6* Pachytene piRNAs

To eliminate production of *pi6* pachytene piRNAs while minimizing the impact on adjacent genes, we used a pair of single-guide RNAs to delete a 227 bp sequence that encompasses the A-MYB-binding site and promoter (Li et al., 2013; Figure 1, S1A, and S1B, and Table S1). For comparison, we created an analogous promoter deletion in *pi17*. We established stable mutant lines (*pi6^em1^*-1, −2, and −3 in Figure S1B) from three founders whose *pi6* promoter deletion sizes range from 219 to 230 bp and differ at their deletion boundaries, reflecting imprecise DNA repair after Cas9 cleavage. All three deletions eliminated *pi6* primary transcripts and mature pachytene piRNAs from both arms of the locus (Figure 1). Because these lines were created using the same pair of sgRNA guides, we refer to all as the *pi6^em1^* allele.

**Figure 1.**
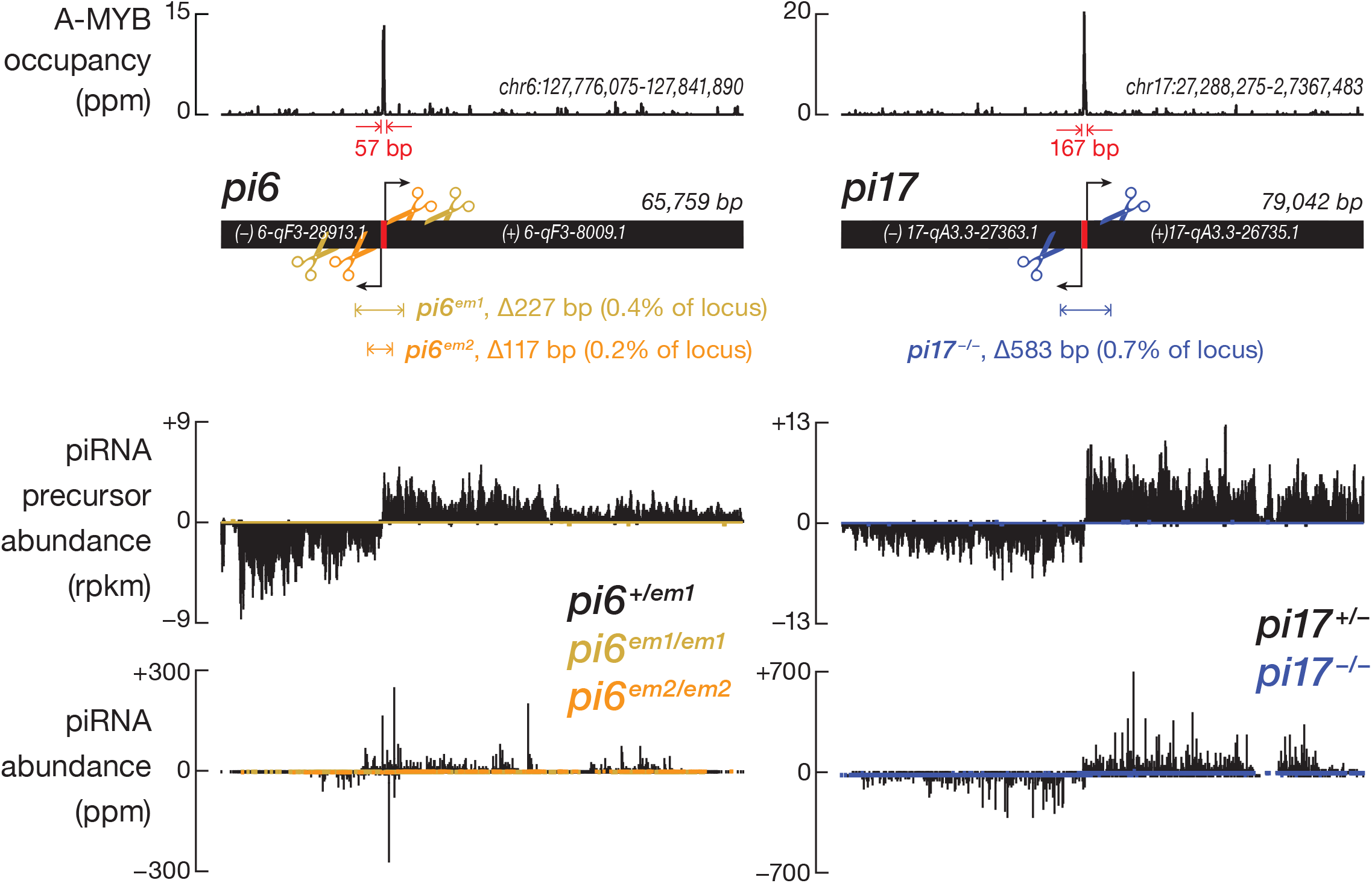
*pi6^em1/em1^*, *pi6^em2/em2^*, and *pi17^−/−^* promoter deletion in mice. Scissors indicate sites targeted by sgRNAs used to guide the Cas9-catalyzed promoter deletions. RNA-seq was used to measure the steady-state abundance of piRNA primary transcripts, and sequencing of NaIO_4_ oxidation-resistant small RNA was used to measure the abundance of mature piRNAs in 17.5 dpp testes. See also Figure S1 and Table S1.

### *pi6* is Required for Male Mouse Fertility

When paired with C57BL/6 females, 2–8 month-old *pi6^em1/em1^* males produced fewer pups compared to their littermates, even at peak reproductive age (Figure 2A and S2A). In six months, C57BL/6 males produced 7 ± 1 (*n* = 5) litters, while *pi6^em1/em1^* males produced 2 ± 2 (*n =* 6) litters. The significantly smaller number of progeny produced by *pi6^em1/em1^* males over their reproductive lifetime does not reflect fewer pups produced in each litter: *pi6^em1/em1^* males sired 5 ± 2 (*n =* 4) pups per litter compared to 6 ± 2 (*n =* 27) for C57BL/6 control males (Figure 2A). Moreover, *pi6^em1/em1^* males regularly produced mating plugs, a sign of coitus, in cohabiting females. Instead, the reduced progeny from *pi6^em1/em1^* males reflects two abnormal aspects of their fertility (Figure 2B). First, 29% of *pi6^em1/em1^* males never produced pups. Second, the mutants that did sire pups did so less frequently. These defects are specific for the loss of *pi6* piRNAs in males because *pi6^+/em1^* heterozygous males and *pi6^em1/em1^* homozygous mutant females showed no discernable phenotype. In contrast, males and females carrying a ∼583-bp promoter deletion in *pi17* were fully fertile, as observed previously for an independent, partial-loss-of-function *pi17* promoter deletion (Homolka et al., 2015), despite loss of primary transcripts and mature piRNAs from both arms of the *pi17* locus (Figure 1).

**Figure 2.**
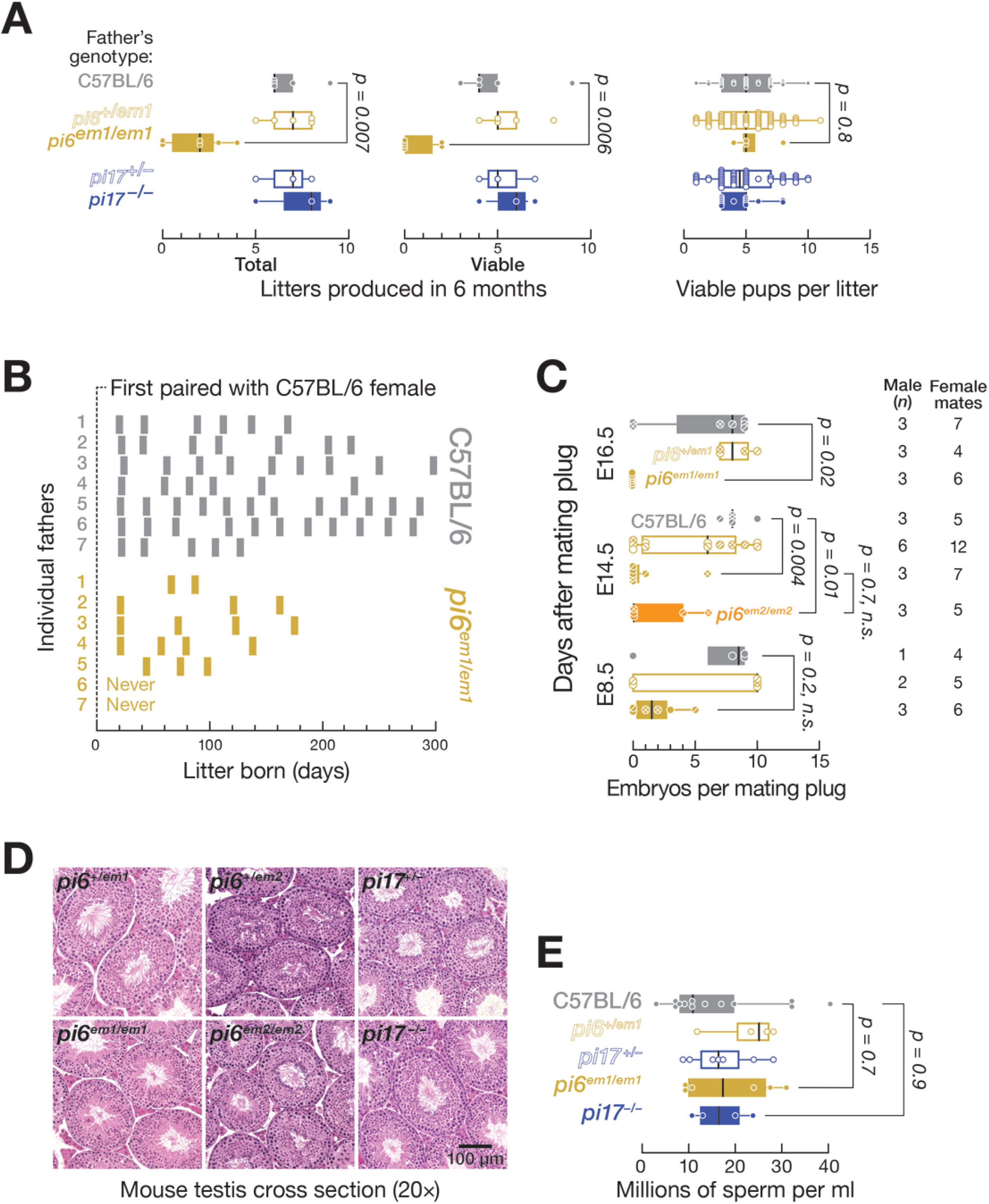
Reduced fertility in *pi6^em1/em1^* males by natural mating. **(A)** Number of litters and pups per litter produced by male mice between 2–8 months of age. **(B)** Frequency and periodicity of litter production. Each bar represents a litter. **(C)** Number of embryos produced by males mated with C57BL/6 females. **(D)** Testis morphology analyzed by hematoxylin and eosin staining. **(E)** Concentration of sperm from the caudal epididymis. In **(A)**, **(C)**, and **(E)**, vertical black lines denote median; whiskers report 75^th^ and 25^th^ percentiles. See also Figure S2.

To test that the reduced fertility of *pi6^em1/em1^* male mice reflects loss of the *pi6* promoter—and not an off-target mutation elsewhere in the genome—we used Cas9 and a second pair of sgRNAs to generate a 117 bp *pi6* promoter deletion, *pi6^em2^* (Figures 1, S1A, and S1C, and Table S1). Like *pi6^em1/em1^* male mice, *pi6^em2/em2^* males produced neither primary *pi6* transcripts nor mature *pi6* piRNAs and showed reduced fertility (Figure S2A). We conclude that *pi6* piRNAs are required for male fertility in C57BL/6 mice.

### *pi6* Mutant Males Produce Fewer Embryos

*pi6* mutant male matings produced fewer fully developed embryos. We examined the embryos produced by natural mating of C57BL/6 females housed with C57BL/6, *pi6^+/em1^*, or *pi6^em1/em1^* males at 8.5, 14.5, or 16.5 days after occurrence of a mating plug. At 8.5 days after mating, C57BL/6 females housed with *pi6^em1/em1^* males carried fewer embryos (2 ± 2, *n =* 3) compared to females paired with *pi6^+/em1^* (6 ± 5, *n* = 2) or C57BL/6 control (7 ± 4, *n =* 1) males (Figure 2C). At 14.5 and 16.5 days post-mating, female mice paired with *pi6^em1/em1^* males had even fewer embryos. Female mice paired with *pi6^em2/em2^* males similarly had fewer embryos at 14.5 days after mating (2 ± 3, *n* = 3). Naturally-born pups sired by *pi6^em1/em1^* and *pi6^em2/em2^* males were rare but healthy, with no obvious abnormalities.

Moreover, *pi6* piRNAs appear to play little if any role in the soma of the developing embryo. *pi6^+/em1^* heterozygous males mated to *pi6^+/em1^* heterozygous females yielded progeny at the expected Mendelian and sex ratios. Moreover, the weight of *pi6^em1/em1^* homozygous pups (28.3 ± 0.6 g, *n =* 6) that developed to adulthood was indistinguishable from their C57BL/6 (26.9 ± 0.3 g, *n =* 6) or heterozygous littermates (28.6 ± 0.3 g, *n =* 8; Figure S2B). We detected no difference in the gross appearance or behavior among these pups.

### *pi6* Mutant Males Produce Mature Spermatozoa

Two-to-four months after birth, both *pi6^+/em1^* and *pi6^em1/em1^* testes weighed ∼15% less than wild-type C57BL/6 testes (Figure S2B). Nonetheless, *pi6^em1/em1^* and *pi6^em2/em2^* testes had normal gross histology, with all expected germ cell types present in seminiferous tubules and sperm clearly visible in the lumen (Figure 2D). The quantity of caudal epididymal sperm produced by *pi6^em1/em1^* mice (19 ± 10 million sperm per ml; *n* = 6) was also comparable to that of their *pi6^+/em1^* (23 ± 7 million sperm/ml; *n =* 4) or C57BL/6 (20 ± 10 million sperm per ml; *n* = 13) littermates (Figure 2E).

Despite the normal quantity of sperm, *pi6^em1/em1^* sperm showed signs of agglutination after 90 min incubation in vitro, compared to C57BL/6 sperm. Moreover, 11 ± 3% (*n* = 4) of *pi6^em1/em1^* caudal epidydimal sperm had abnormal head morphology (Figure S2C), compared to 2 ± 1% (*n* = 5) of wild-type sperm (*p = 0.02*). Defects in germ cell chromosomal synapsis, triggering errors in gene expression, have been linked to abnormal sperm head shape (Wong et al., 2008; de Boer et al., 2015). In fact, 22 ± 7 percent (*n* = 4) of *pi6^em1/em1^* pachytene spermatocytes had unsynapsed sex chromosomes or incompletely synapsed autosomal chromosomes, compared to 7 ± 3 percent (*n =* 4) for C57BL/6 (Figure S2D and S2E).

### *pi6* Mutant Sperm Fail to Fertilize Wild-type Eggs

Because *pi6*^−/−^ males are ineffectual at siring offspring, we used in vitro fertilization (IVF) to distinguish between defects in mating behavior and sperm function, incubating sperm from C57BL/6, *pi6^+/em1^*, or *pi6^em1/em1^* males with wild-type oocytes and scoring for the presence of both male and female pronuclei and the subsequent development of the resulting bi-pronuclear zygotes into two-cell embryos 24 h later (Figure 3A). The majority of oocytes incubated with sperm from C57BL/6 (86 ± 17%; 774 total oocytes; *n* = 6) or *pi6^+/em1^* (60 ± 35%; 412 total oocytes; *n* = 3) males developed into two-cell embryos. By contrast, only 7 ± 5% (12-fold decrease compared to C57BL/6; Cohen’ *d* = 6.3; 1,026 total oocytes; *n* = 7) of oocytes incubated with *pi6^em1/em1^* sperm reached the two-cell stage. Similarly, no oocytes incubated with *pi6^em2/em2^* sperm developed into two-cell embryos by 24 h. The majority of these oocytes remained undivided, and few contained a male pronucleus, suggesting that *pi6^em1/em1^* and *pi6^em2/em2^* sperm are defective in fertilization.

**Figure 3.**
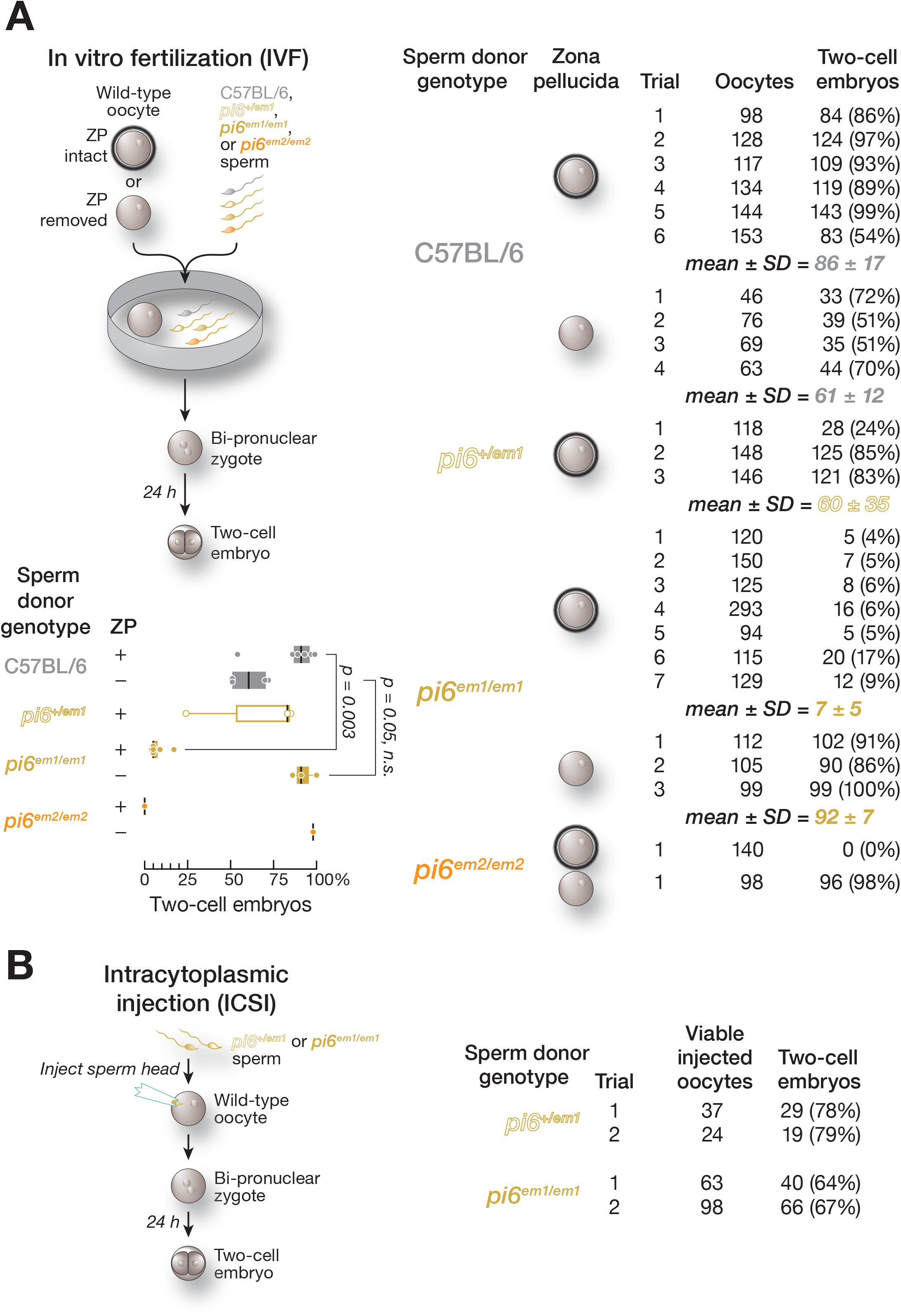
Fertilization defects of *pi6^em1/em1^* and *pi6^em2/em2^* sperm revealed by IVF and ICSI. **(A)** Sperm function analyzed by in vitro fertilization (IVF) using oocytes with or without zona pellucida. Vertical black lines denote median; whiskers report the 75^th^ and 25^th^ percentiles. **(B)** Sperm function analyzed by intracytoplasmic sperm injection (ICSI). See also Figure S3

### *pi6^em1/em1^* Sperm Nuclei Support Fertilization

The best studied piRNA function is transposon silencing, and mouse *pi2* has been proposed to be involved in LINE1 element silencing, although *pi2* mutant males are fertile (Xu et al., 2008). Moreover, LINE1 transcript abundance increases in mice bearing inactivating mutations in the catalytic site of MIWI (Reuter et al., 2011). Transposon activation can produce DNA damage, and genomic integrity is critical for fertilization (Ahmadi and Ng, 1999; Morris et al., 2002; Bourc’his and Bestor, 2004; Lewis and Aitken, 2005). However, pachytene piRNAs are depleted of repetitive sequences in contrast to other types of piRNA-producing genomic loci (Figure S3A; Aravin et al., 2006; Girard et al., 2006; Gainetdinov et al., 2018).

We asked whether the defect in fertilization by *pi6* mutant sperm might reflect DNA damage or epigenetic dysregulation of the sperm genome. *pi6^+/em1^ or pi6^em1/em1^* sperm heads were individually injected into the cytoplasm of wild-type oocytes (intracytoplasmic sperm injection, ICSI; Figure 3B), bypassing the requirement for sperm motility, acrosome reaction, egg binding, or sperm-egg membrane fusion (Kuretake et al., 1996). *pi6^em1/em1^* sperm heads delivered by ICSI fertilized the oocyte at a rate similar to that of *pi6^+/em1^* sperm: 66% of oocytes (161 total viable oocytes from two separate trials) injected with homozygous mutant *pi6^em1/em1^* sperm heads reached the two-cell stage, compared to 79% for *pi6^+/em1^* (61 total viable oocytes from two separate trials). Thus, most *pi6^em1/em1^* nuclei are capable of fertilization.

The steady-state abundance of transposon RNA in *pi6^em1/em1^* and *pi6^em2/em2^* testicular germ cells further supports the view that the fertilization defect caused by loss of *pi6* piRNAs does not reflect a failure to silence transposons. We used RNA-seq to measure the abundance of RNA from 1,007 transposons in four distinct purified germ cell types: pachytene spermatocytes (4C), diplotene spermatocytes (4C), secondary spermatocytes (2C), and spermatids (1C). *pi6* piRNAs compose 5.5% of all spermatocyte pachytene piRNAs (943,758 molecules per diplotene spermatocyte; Figure S3B), yet when *pi6* piRNAs were eliminated in *pi6^em1/em1^* or *pi6^em2/em2^* mice, we found no significant changes in steady-state RNA abundance (i.e., an increase or decrease >2-fold and FDR <0.05) for any transposon family compared to C57BL/6 germ cells (Figure S3C). We also note that, as in C57BL/6 wild-type, ɣH2AX expression was confined to the sex body in pachytene spermatocytes in *pi6^em1/em1^* testis, indicating an absence of DNA damage (data not shown). Together with the rescue of the fertilization defects of *pi6^em1/em1^* sperm by ICSI, these data suggest that transposon silencing is unlikely to be the essential biological function of *pi6* piRNAs.

### *pi6* Mutant Sperm Struggle to Penetrate the Zona Pellucida

Mammalian spermatozoa stored in the epididymis are immotile and dormant. Sperm capacitate, i.e., resume maturation, only upon entering the female reproductive tract (de Lamirande et al., 1997). Upon capacitation, sperm become capable of undergoing the acrosome reaction, which is required to bind and penetrate the outer oocyte glycoprotein layer, the zona pellucida (Florman and Storey, 1982; de Lamirande et al., 1997; Jin et al., 2011). To test whether the defect in fertilization by *pi6* mutant sperm was due to impaired binding to or penetration of the zona pellucida, we compared IVF using unmanipulated oocytes to oocytes with the zona pellucida removed (Figure 3A). Strikingly, removing the zona pellucida fully rescued the fertilization rate of *pi6 ^em1/em1^* sperm: 92 ± 7% (316 total oocytes; *n* = 3) of zona pellucida-free oocytes incubated with *pi6^em1/em1^* sperm reached the two-cell stage after 24 h, compared to those with intact zona pellucida (7 ± 5%; 1,026 total oocytes; *n* = 3; Figure 3A). Similarly, 98% of zona pellucida-free oocytes incubated with *pi6^em2/em2^* sperm developed into two-cell embryos after 24 h (98 total oocytes; *n* =1), in contrast to 0% of those with intact zona pellucida (140 total oocytes; *n* = 1).

### Impaired Capacitation in *pi6* Mutant Sperm

One hallmark of sperm capacitation is a switch to “hyperactivated motility,” a swimming pattern characterized by a high amplitude and non-symmetric beating of the flagellum that facilitates penetration of the zona pellucida (Suarez et al., 1991; Stauss et al., 1995; Quill et al., 2003; Qi et al., 2007). To assess *pi6* mutant sperm capacitation, we measured the motility of freshly extracted caudal epididymal sperm from *pi6^em1/em1^*, *pi6^em2/em2^,* or C57BL/6 mice using computer-assisted sperm analysis (CASA; Figure 4A; Mortimer, 2000). After 90 min incubation under capacitation-promoting conditions, *pi6^em1/em1^* and *pi6^em/em2^* sperm populations had reduced path and progressive velocity, measures of sperm motility, compared to control sperm (Figure 4B and 4C). In fact, 10 min after sperm extraction, most *pi6^em1/em1^* sperm moved more slowly than C57BL/6 control sperm (Movies S1 and S2), and as the sperm were incubated in capacitating conditions, motility of the mutant sperm motility continued to declined more rapidly than that of control (Movies S3–S10). By 4 h, most *pi6^em1/em1^* sperm only moved in place and showed signs of agglutination (Movies S8 and S10).

**Figure 4.**
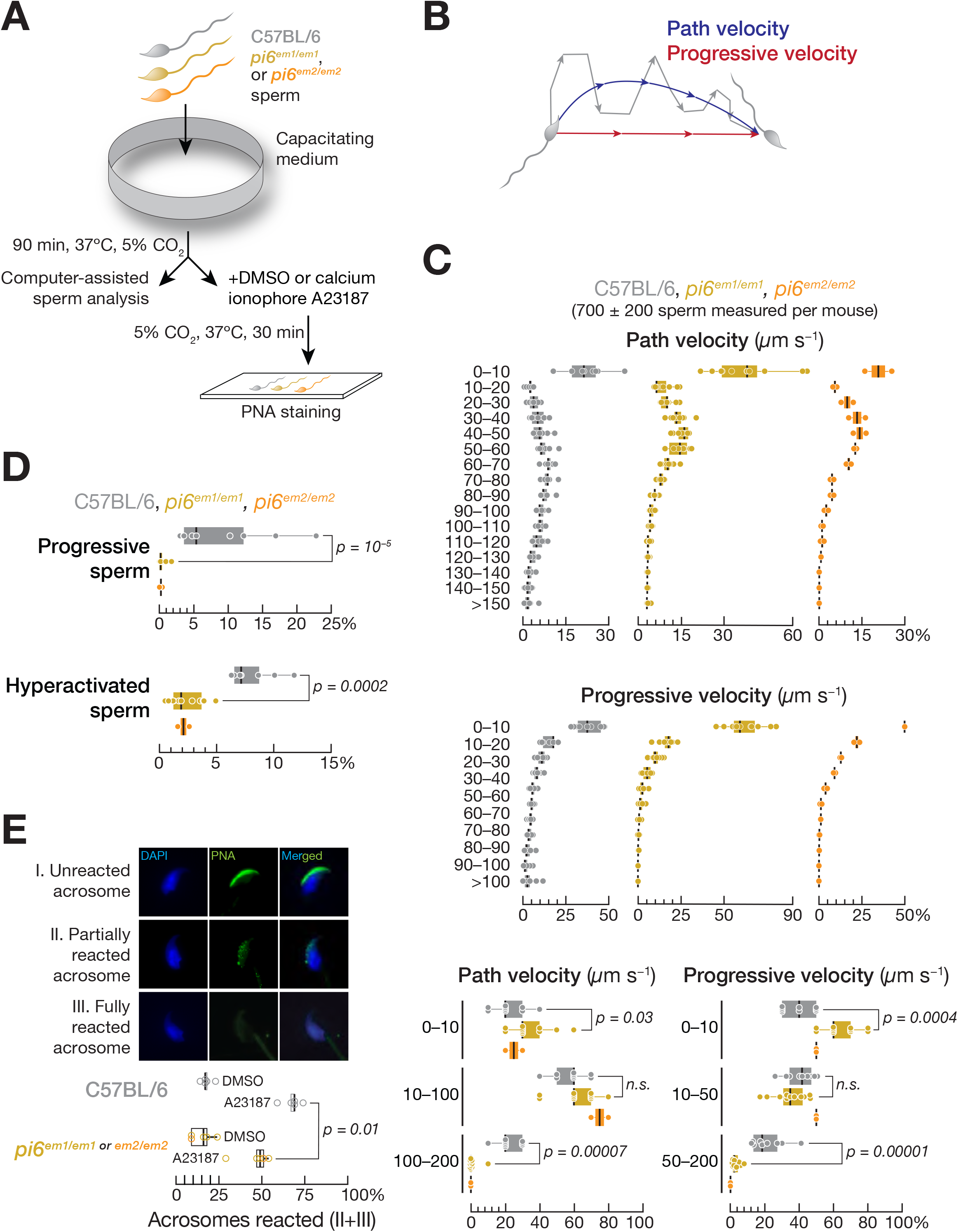
Impaired sperm capacitation in *pi6* mutant sperm. **(A)** Strategy to measure sperm motility and acrosome reaction triggered with Ca^2+^ ionophore A23187. **(B)** Definition of path and progressive velocities. **(C)** Distribution of path and progressive velocities for sperm from C57BL/6 (*n* = 9), *pi6^em1/em1^* (*n* = 11), and *pi6^em2/em2^* (*n* = 2). Top: 10 µm/sec bins; bottom: bins correspond to immotile or slow, intermediate, and vigorous motility. **(D)** Distribution of progressive and hyperactivated sperm from C57BL/6, *pi6^em1/em1^*, and *pi6^em2/em2^* mice determined by CASAnova. **(E)** (Top panel) Acrosome status of representative wild-type caudal epididymal spermatozoa. Green, peanut agglutinin to detect the acrosome; blue, DAPI to detect DNA. (Bottom panel) Acrosome reaction rates for wild-type and *pi6* mutant sperm. The results using *pi6^em1/em1^* and *pi6^em2/em2^* sperm for acrosome reaction were combined as indicated. In **(C)**, **(D)**, and **(E)**, vertical black lines denote median; whiskers report 75^th^ and 25^th^ percentiles. See also Movies S1–S10.

To more rigorously evaluate progressive motility and hyperactivation, we used CASAnova, an unsupervised machine learning tool (Goodson et al., 2011), to analyze caudal epididymal sperm from *pi6^em1/em1^*, *pi6^em2/em2^,* and C57BL/6 control mice. After 90 min in capacitating conditions, CASAnova identified just 0.3 ± 0.5% of *pi6^em1/em1^* (*n* = 11) and 0.2 ± 0.3% of *pi6^em2/em2^* (*n* = 2) sperm as progressive, compared to 9 ± 7% for C57BL/6 (*n* = 9; Figure 4D). Similarly, only 2 ± 1% of *pi6^em1/em1^* or *pi6^em2/em2^* sperm displayed hyperactivated motility, compared to 8 ± 2% for the control (Figure 4D), a percentage typical for the C57BL/6 mouse strain (Goodson et al., 2011).

Ex vivo, the acrosome reaction occurs spontaneously in some sperm and can also be triggered by the Ca^2+^ ionophore A23187 (Talbot et al., 1976), which results in an acrosome reaction visually indistinguishable from that triggered by natural ligands such as progesterone (Osman et al., 1989) or ZP3 (Arnoult et al., 1996), while bypassing signaling pathways essential for acrosome reaction in vivo (Tateno et al., 2013; Figure 4A and 4E). While the spontaneous acrosome reaction rates for C57BL/6 (18 ± 3%; *n* = 5) and *pi6* mutant sperm were similar (15 ± 6%; *n* = 5), acrosome reaction triggered by ionophore-induced Ca^2+^ influx (i.e., ionophore-induced minus spontaneous) differed between the two genotypes: 46 ± 10% (31 ± 12%) of *pi6* mutant sperm (*n* = 5) underwent partial or complete reaction, compared to 68 ± 6% (50 ± 7; *n* = 5) for C57BL/6 (*p* = 0.01; Cohen’s *d* = 2.78; Figure 4E). Our data suggest that *pi6* mutant sperm less effectively undergo an acrosome reaction triggered by ionophore-induced Ca^2+^ influx, a defect expected to impair binding and penetrating the zona pellucida. Together, our data indicate that insufficient capacitation is responsible for the poor fertilization capability of *pi6* mutant sperm.

### Potential Role of Paternal *pi6* piRNAs in Embryo Development

Even when *pi6* mutant sperm successfully fertilize an oocyte, the resulting heterozygous embryos are less likely to complete gestation. Two-cell embryos generated by IVF using sperm from *pi6^+/em1^*, *pi6^em1/em1^*, or C57BL/6 control mice were transferred to wild-type surrogate mothers (Figure 5A). At least half the embryos from *pi6^+/em1^* (50 ± 10%; 23 ± 4 embryos per female; *n* = 3) or C57BL/6 control sperm (70 ± 10%; 21 ± 3 embryos per female; *n* = 3) developed to term (Figure 5B), a rate typical for this genetic background (González-Jara et al., 2017). By contrast, only 20 ± 20% of two-cell embryos from *pi6^em1/em1^* sperm developed to term (*n* = 6).

**Figure 5.**
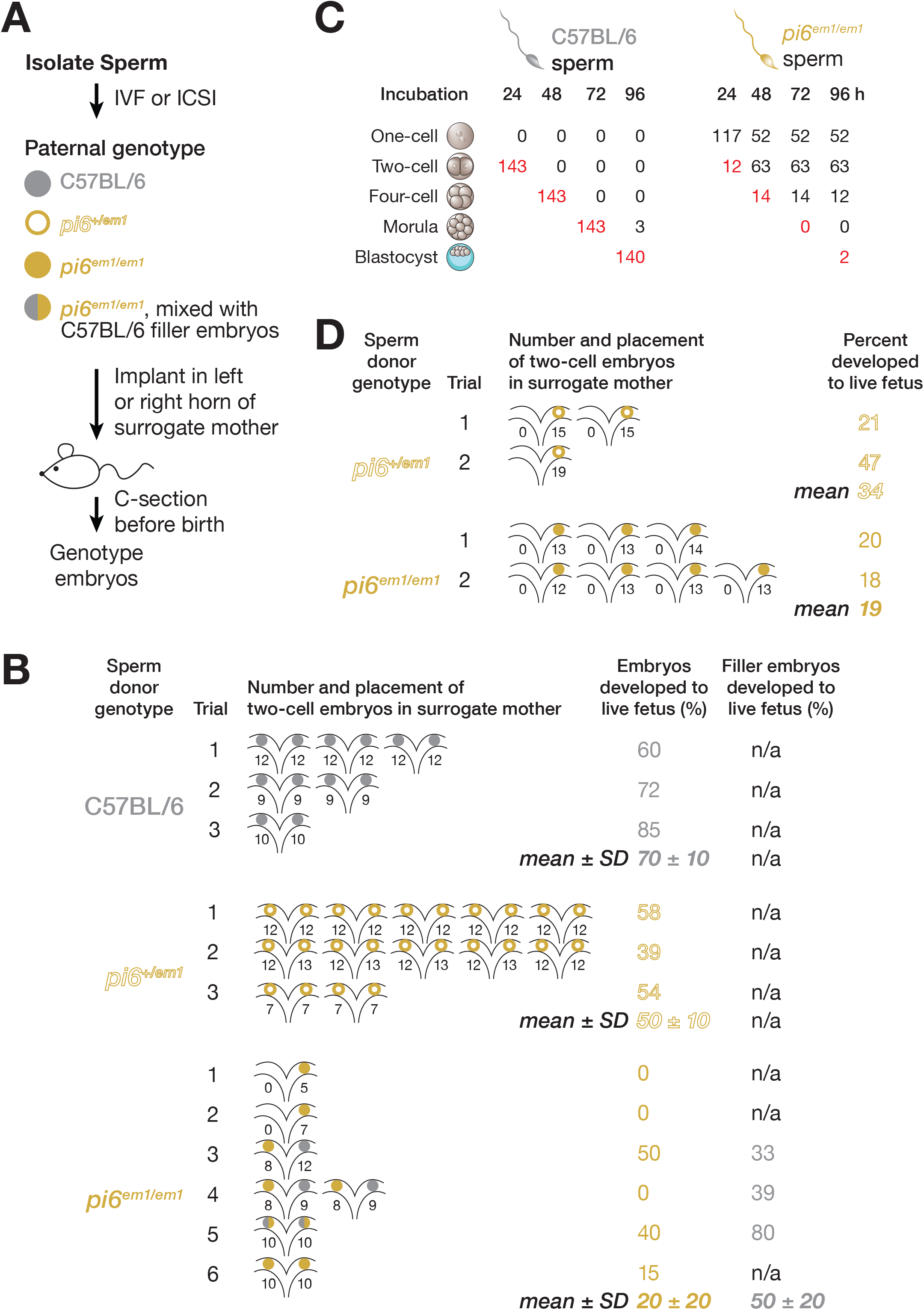
Embryos derived from *pi6^em1/em1^* sperm fail to develop. **(A)** Strategy for surgical transfer of fertilized two-cell embryos to surrogate mothers. **(B)** Percentages of IVF-derived two-cell embryos that developed to term. Each uterine cartoon represents one surrogate mother; colored circles depict embryos. The number of embryos transferred to each side of the oviduct is indicated. **(C)** Development of IVF-derived embryos. Red, number of embryos that developed to the stage appropriate for the elapsed time after fertilization. **(D)** Percentages of ICSI-derived two-cell embryos that developed to term. See also Figure S4

The low number of fertilized two-cell embryos produced in IVF using *pi6^em1/em1^* sperm precluded transferring the standard number of embryos to surrogate mothers. For example, in two IVF experiments using *pi6^em1/em1^* sperm, only 5 or 7 embryos could be transferred; the surrogate females failed to become pregnant (Figure 5B and S4A, Trials 1 and 2). In theory, this result might suggest a paternal role for *pi6*. A more mundane explanation is that the low number of embryos transferred reduced the yield of live fetuses, as reported previously (McLaren, 1955; Johnson et al., 1996; González-Jara et al., 2017). We conducted additional experiments to distinguish between these two possibilities. Oocytes were again fertilized by IVF with *pi6^em1/em1^* or C57BL/6 control sperm, and two-cell embryos transferred to surrogate females, but matching the number of embryos transferred to each surrogate for the two sperm genotypes. We used two strategies. First, similar numbers of embryos derived from *pi6^em1/em1^* sperm and wild-type “filler” embryos derived from control sperm were transferred to separate oviducts in the same females to make up the total numbers of transferred embryos (Figure 5B, Trials 3 and 4). Again, fewer embryos developed to term for *pi6^em1/em1^* (17%) compared to control sperm (37%). Second, embryos derived from *pi6^em1/em1^* sperm and wild-type filler embryos were mixed before transfer and then equal numbers of embryos, selected randomly, were implanted in each oviduct (Figure 5B, Trial 5). Pups isolated by cesarean section 18.5 days after transfer were genotyped by PCR. In this experiment, only 40% of embryos derived from *pi6^em1/em1^* sperm developed to term, compared to 80% of wild-type filler embryos. Finally, in one experiment (Trial 6) where we obtained sufficient numbers of embryos derived from *pi6^em1/em1^* sperm, 10 *pi6^em1/em1^*-derived two-cell embryos were transferred to each oviduct of the surrogate female. Just 15% of the *pi6^em1/em1^*-derived embryos developed to term, compared to 85% for C57BL/6. Together, these data suggest a direct or indirect paternal role for *pi6* piRNAs in the embryo.

We also monitored pre-implantation development in vitro for up to 96 h, a period during which the one-cell embryo develops into a blastocyst. Of the oocytes incubated with *pi6^em1/em1^* sperm, 40% remained undivided without evidence of a male pronucleus, presumably because they were not fertilized. Among the remaining 60% oocytes that progressed to at least the two-cell stage, indicating successful fertilization by *pi6^em1/em1^* sperm, 82% showed delayed development, requiring 48 h to reach the two-cell stage. None of these developed further. Just 3% of oocytes fertilized by *pi6^em1/em1^* sperm progressed to the blastocyst stage by 96 h, compared to 98% for C57BL/6 sperm (Figure 5C).

Further support for the idea that paternal *pi6* piRNAs play a role in embryogenesis or embryonic viability comes from transfer of embryos generated by ICSI (Figure 5D). ICSI with *pi6^em1/em1^* or *pi6^+/em1^* sperm yielded comparable normal numbers of fertilized oocytes (Figure 3B), so no wild-type filler embryos were used; all embryos were transferred into a single oviduct of the surrogate female. In two independent experiments in which embryos generated by ICSI were transferred to surrogate mothers, only 19% of two-cell embryos derived from *pi6^em1/em1^* sperm heads developed to term, compared to 34% for embryos fertilized with *pi6^+/em1^* (Figure 5C). Only four of seven (57%) surrogate mothers carrying embryos derived from *pi6^em1/em1^* sperm became pregnant. All three surrogate mothers receiving embryos derived from *pi6^+/em1^* sperm became pregnant (Figure S4B).

We note that the live fetuses generated using *pi6^em1/em1^* sperm in IVF or sperm heads in ICSI, like those produced by natural mating using *pi6^em1/em1^* males, showed no obvious morphological abnormalities and grew to adulthood normally when fostered by host mothers. We conclude that paternal *pi6* piRNAs play a direct or indirect role in early embryogenesis.

### *pi6* Promoter Deletion Does Not Cause Large-Scale Changes in the Expression of Neighboring Genes

In theory, disruption of *pi6* could influence flanking gene expression, confounding transcriptome analysis. However, we find no evidence for coincidental changes in the expression of the genes flanking *pi6*. In *pi6* mutant pachytene spermatocytes, diplotene spermatocytes, and secondary spermatocytes, no gene on chromosome 6 is affected except for *pi6* itself. In spermatids, the steady-state mRNA abundance of *Atp6v1e1*, which lies 7 Mb upstream of *pi6*, more than doubled in *pi6* mutants (2.1- and 2.3-fold increase in *pi6^em1/em1^* and *pi6^em2/em2^* spermatids, respectively), but expression of intervening genes was unaltered. Among the genes between *pi6* and *Apt6v1e1*, 19 have mRNAs with abundance >10 molecules per cell in spermatids; none were affected by loss of *pi6* transcription. The widespread preservation of normal mRNA abundance for genes on chromosome 6 strongly argues that loss of *pi6* transcription has little or no effect on the chromatin structure of neighboring genes.

### Changes in Spermatocyte and Spermatid mRNA Abundance Accompany Loss of *pi6* piRNAs

Pachytene piRNAs repress their RNA targets at least in part by an siRNA-like cleavage mechanism. Mice bearing mutations that selectively inactivate the endonuclease activity of MIWI are phenotypically indistinguishable from those lacking MIWI altogether (Deng and Lin, 2002; Reuter et al., 2011). Moreover, ectopic expression in mice of the largest human piRNA-producing locus triggers cleavage and degradation of mouse *Dpy19l2* mRNA, causing male sterility (Goh et al., 2015). Therefore, to begin to identify direct targets of *pi6* piRNAs, we first used RNA-seq to measure steady-state RNA abundance in pachytene spermatocytes, diplotene spermatocytes, secondary spermatocytes, and spermatids purified from *pi6^em1/em1^*, *pi6^em2/em2^*, and C57BL/6 adult testis (Figure 6A).

**Figure 6.**
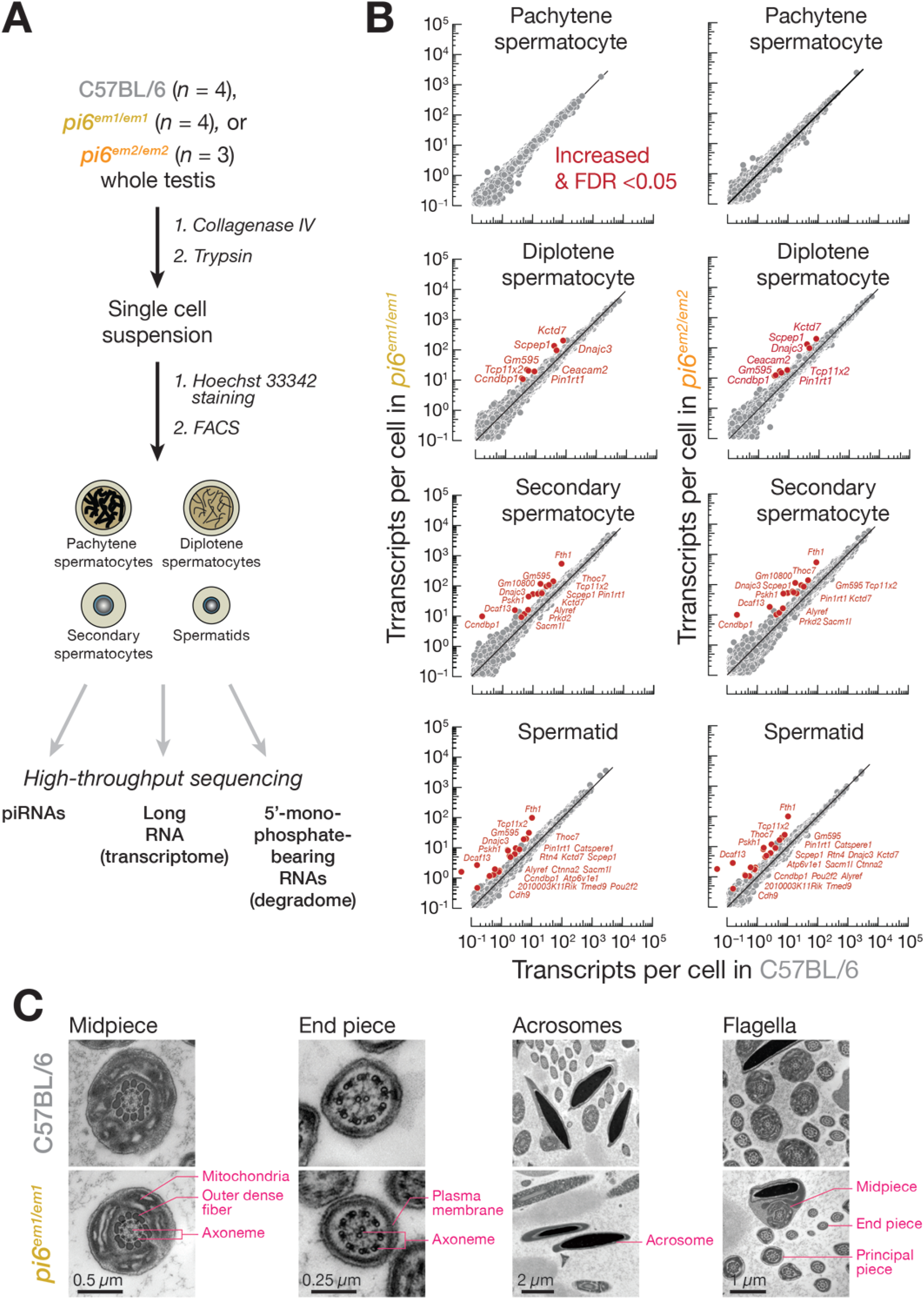
mRNAs encoding proteins required for sperm capacitation and zona pellucida-binding are direct targets of *pi6* piRNAs. **(A)** Strategy for purifying specific male germ cell types. **(B)** Scatter plots of steady-state transcript abundance in sorted testicular germ cells. Each dot represents the mean abundance of an mRNA measured using four (wild-type and *pi6^em1/em1^* cells) or three (*pi6^em2/em2^* cells) biologically independent samples. Differentially expressed transcripts (>2 fold-change and <0.05 FDR) are indicated. **(C)** Ultrastructure of caudal epididymal sperm flagella and acrosomes from mice of indicated genotypes by transmission electron microscopy. See also Figures S5 and S6, and Tables S2, S3, and S4.

The steady-state abundance of the RNA targets of *pi6* piRNA-guided cleavage are predicted to increase in *pi6* mutants. We searched for transcripts whose steady-state abundance increased in both *pi6^em1/em1^* (*n* = 4) and *pi6^em2/em2^* (*n* = 3) cells compared to C57BL/6 controls (*n* = 4; Figure 6B and Table S2). *pi6^em1^* and *pi6^em2^* deletions increased the abundance of 8 diplotene spermatocyte mRNAs, 15 secondary spermatocyte mRNAs, and 21 spermatid mRNAs. Although *pi6* piRNAs first begin to accumulate in pachytene spermatocytes (Figure S3B), the abundance of no pachytene spermatocyte mRNA changed significantly (FDR <0.05) in both *pi6^em1/em1^* and *pi6^em2/em2^* mice, suggesting that *pi6* piRNAs do not accumulate to functional levels until the diplotene phase of meiosis. In total, loss of *pi6* piRNAs more than doubled the mRNA level of 24 genes in at least one spermatogenic cell type, 13 (54%) of which remained significantly altered in subsequent stages.

### Genes Essential for Sperm Functions are Regulated by *pi6* piRNAs

Among the 24 genes with increased mRNA abundance in *pi6* mutant cells, *Atp6v1e1* and *Catspere1* encode proteins required for sperm function (Table S3). ATP6V1E1, the testis-specific, catalytic subunit of the vacuolar-type H^+^ ATPase, resides in the inner and outer-membranes of the acrosome and acts to acidify the acrosome, stabilizing enzymes required for sperm to penetrate the oocyte zona pellucida (Huang et al., 1985; Sun-Wada et al., 2002). Although *Atp6v1e1* overexpression has not been examined in mouse spermatogenesis, overexpression of *Atp6v1e1* in somatic tissues is associated with cancer (Son et al., 2016). In *pi6^em1/em1^* and *pi6^em2/em2^* spermatids, *Atp6v1e1* mRNA expression increased by 2.1-(FDR = 2 × 10^−10^) and 2.3-fold (FDR = 5 × 10^−7^), respectively (Figure 6 and Table S2). *CATSPERE1* is one of the nine subunits of the sperm-specific CatSper calcium channel, which resides in the flagellar membrane and is required for the transition to hyperactivated motility during capacitation (Chung et al., 2017). In humans, a homozygous in-frame deletion of *Catspere1* prevents sperm from fertilizing oocytes, resulting in male infertility (Brown et al., 2018).

In addition to the well-defined sperm functions of *Atp6v1e1* and *Catspere1*, *Ceacam2*, *Pou2f2*/*Oct2*, and *Tcp11x2*, genes whose mRNA abundance increases in *pi6* mutants, have been proposed to function in spermatogenesis. *Ceacam2* encodes CEACAM2-L, a testis-specific isoform of the carcinoembryonic antigen-related cell adhesion molecule (CEACAM) family of proteins. CEACAM2-L appears in elongated spermatids and becomes undetectable in epididymal sperm, suggesting a role as a cell surface, testicular cell adhesion factor (Salaheldeen et al., 2012). *Pou2f2* encodes an transcription factor that binds DNA cooperatively with other POU domain-containing proteins (Verrijzer et al., 1992; Lim et al., 2011); *Pou2f2* is normally expressed in pre-meiotic, type-A spermatogonia, but not in gonocytes, meiotic germ cells, or post-meiotic germ cells (Lim et al., 2011). The function of *Tcp11x2*, which encodes an X-linked T-complex 11 protein, is suggested by its well-characterized paralog, TCP11, a receptor for fertilization-promoting peptides that facilitate sperm capacitation (Fraser et al., 1997; Adeoya-Osiguwa et al., 1998; Ma et al., 2002). Seventeen additional genes whose mRNA abundance increased in *pi6* mutants have reported functions only in somatic cells; the functions of two (*Gm595* and *2010003K11Rik*) are unknown (Table S3).

### *pi6*-Regulated Genes Regulate Related Cellular Processes

Loss-of-function phenotypes have been reported for 15 of the 24 *pi6* piRNA repressed genes. Knockout of nine genes (*Atp6v1e1*, *Rtn4*, *Ctnna2*, *Pskh1*, *Sacm1l*, *Dnajc3*, *Pouf2f2*, *Dcaf13*, *Fth1*) leads to embryonic or neonatal lethality or premature death in mice (Table S3). Two-thirds of the 24 genes encode subunits of multi-protein complexes with known functions. Overexpression of individual subunits can disrupt the function of a complex by a variety of mechanisms (Prelich, 2012). Therefore, although in vivo knockout studies have not been reported for many *pi6*-regulated genes, their functions can be inferred from that of the larger complex in which they reside. For example, loss of THOC1 or THOC5 in the TREX complex, which also contains THOC7 and ALYREF/THOC4, or of TMED2, which forms a complex with TMED9, causes embryonic lethality (Wang et al., 2006; Jerome-Majewska et al., 2010; Mancini et al., 2010). For 12 of the *pi6*-regulated genes, knockout mutants die before puberty, preventing assessment of a role in male fertility using existing alleles. Heterozygous mutants of one of these homozygous lethal genes, *Sacm1l*, have abnormal testis or epididymis morphology, suggesting that *Sacm1l* is important for spermatogenesis (Koscielny et al., 2014).

In addition to ATP6V1E1 and CATSPERE1, eight *pi6* piRNA-regulated genes encode integral membrane proteins or regulators of membrane biogenesis or function (*Cdh9*, *Ceacam2, Dnajc3*, *Kctd7*, *Rtn4*/*Nogo*, *Sacm1l*, *Tmed9*, and |*Piro-1*/Gm10800; Nemoto et al., 2000; Shimoyama et al., 2000; Ladiges et al., 2005; Voeltz et al., 2006; Liu et al., 2008; Azizieh et al., 2011; Jozsef et al., 2014; Oh et al., 2015; Ghanem et al., 2016; Dvela-Levitt et al., 2019). These proteins could play a role in acrosome function. Another 8 *pi6*-regulated genes function in pathways that control or respond to intracellular ion levels. *Catspere1*, *Cdh9*, *Ceacam2*, *Prkd2*, *Pskh1*, and *Rtn4* encode proteins that regulate or respond to intracellular Ca^2+^ concentration (Edlund and Obrink, 1993; Hunter et al., 1996; Brede et al., 2000; Shimoyama et al., 2000; Shapiro and Weis, 2009; Jozsef et al., 2014; Ghanem et al., 2016; Chung et al., 2017; Xiao et al., 2018). The protein products of *Atp6v1e1* and *Fth1* are required for intracellular H^+^ and Fe^3+^ homeostasis, respectively (Ferreira et al., 2000; Sun-Wada et al., 2002; Pietrement et al., 2006). Thus, altered ion homeostasis may explain at least some of the defects of *pi6* sperm.

Three *pi6*-regulated genes encode proteins that function in proteasomal protein degradation (KCTD7, DCAF13, DNAJC3). Four mediate cell-cell adhesion (CDH9, CTNNA2 [αN-Catenin], CEACAM2, PRKD2). Two, ALYREF/THOC4 and THOC7, act in mRNA processing or export as components of the TREX complex, which couples mRNA transcription, splicing, and nuclear export (Strässer et al., 2002; Masuda et al., 2005; Chi et al., 2013). Together, the known and inferred functions of *pi6*-regulated genes suggest that the mechanism underlying *pi6* sperm defects reflects dysregulated ion homeostasis rather than aberrant sperm flagellar structure. Consistent with this, transmission electron microscopy detected no architectural abnormalities in the *pi6^em1/em1^* sperm flagellum or acrosome (Figure 6C).

### *pi6* piRNAs Direct Cleavage of Their mRNA Targets

All known catalytically active Argonaute proteins cleave their targets at the phosphodiester bond linking nucleotides t10 and t11, the bases paired to guide nucleotides g10 and g11. Target cleavage generates 5′ product bearing a 3′ hydroxyl terminus and a 3′ product beginning with a 5′ monophosphate. Thus, cleaved targets can be identified by high-throughput sequencing methods designed to capture long RNAs bearing a 5′ monophosphate group (degradome-seq) coupled with computational identification of piRNAs capable of directing production of the putative 3′ cleavage products.

We performed small RNA-seq to define the piRNA repertoire and degradome-seq of C57BL/6 and *pi6^em1/em1^* germ cells to identify candidate *pi6* piRNA-directed target cleavage products. Because the specific rules for piRNA-guided target cleavage are poorly defined, we identified target candidates by first requiring g2–g7 seed complementarity between a *pi6* piRNA and a cleaved RNA fragment. Then, we searched for seed-matched transcripts with a cleavage product whose 5′ end overlapped 10 nt with the piRNA (Figure S5A). Finally, we compared both the steady-state (RNA-seq) and cleaved fragment (degradome-seq) abundance of target candidate RNAs in wild-type and *pi6* mutant germ cells. These criteria identified *pi6* piRNA-dependent cleavage sites in six mRNAs whose abundance increased in *pi6* mutants: *Alyref*, *Catspere1*, *Dnajc3*, *Fth1*, *Kctd7*, and *Scpep1* (Figure S5B and Table S4). For example, a target cleavage site in *Catspere1* mRNA exon 11 is complementary to nucleotides g2–g7 and to an additional 11 nucleotides between g8 and g21 of a *pi6* piRNA present at 744 molecules per cell in diplotene spermatocytes (*n* = 3). In *pi6^em1/em1^* pachytene, diplotene, and secondary spermatocytes, we could no longer detect the 5′ monophosphorylated 3′ product of *Catspere1* mRNA cleavage, and the steady-state abundance of *Catspere1* mRNA increased 2.1-(FDR = 5 × 10^−18^) and 2.2-(FDR = 6 × 10^−8^) times in *pi6^em1/em1^* and *pi6^em2/em2^* spermatids, respectively (Figure 6B and Table S2). We detected similar *pi6*-dependent cleavage sites in *Alyref* (one open-reading frame [ORF] cleavage site in exon 5), *Dnajc3* (one ORF cleavage site in exon 5), *Fth1* (one ORF cleavage site in exon 2; one ORF cleavage site in exon 4), *Kctd7* (two 3′ UTR cleavage sites in exon 4), and *Scpep1* (one ORF cleavage site in exon 4).

To validate targets predicted to be cleaved by *pi6* piRNAs, we used qRT-PCR to measure the abundance of full length mRNA in wild-type and *pi6^em1/em1^* spermatids using primers spanning the putative cleavage site (Figure S5; Figure S6). Consistent with our RNA-seq data, all six *pi6* piRNA target mRNAs increased >3-fold in mutant spermatids (*p* ≤ 0.006, *n* = 3; Figure S5A). Together, our results identify mRNAs directly regulated by *pi6* piRNAs via siRNA-like target cleavage.

### *pi6* piRNAs Reciprocally Facilitate Biogenesis of piRNAs from Other Loci

Because piRNA-directed cleavage of piRNA precursor transcripts generates 5′ monophosphorylated pre-pre-piRNAs, piRNAs play a central role in the initiation of piRNA production (Brennecke et al., 2007; Gunawardane et al., 2007; Han et al., 2015a; Mohn et al., 2015; Gainetdinov et al., 2018). Intriguingly, loss of *pi6* piRNAs specifically decreased the abundance of piRNAs from two pachytene piRNA-producing loci on chromosome 10, but not from any other loci, including the major piRNA loci *pi2*, *pi7*, *pi9*, or *pi17* (Figure 7A). Moreover, loss of *pi6* piRNAs had no significant effect (>2 fold-change and <0.05 FDR) on the abundance of mRNAs encoding piRNA pathway proteins or piRNAs derived from major piRNA-producing loci (Figure S3B and S6B; Table S5). At their peak expression in wild-type diplotene spermatocytes, *pi10-qC2-545.1* (*chr10: 94,673,712–94,688,442*) produces 0.15 ± 0.01% (26,192 molecules per diplotene spermatocyte, *n* = 3) of pachytene piRNAs, while *pi10-qA3-143.1* (*chr10: 20,310,505–20,312,312*) produces 0.0259 ± 0.0003% (4,470 molecules per diplotene spermatocyte, *n* = 3). In *pi6* mutants, *pi10-qC2-545.1* piRNAs decreased 2.5-fold in pachytene spermatocytes (*p* = 0.005) and ≥3-fold in diplotene spermatocytes, secondary spermatocytes, and spermatids (*p* = 0.0001). *pi10-qA3-143.1* piRNAs declined 1.7-fold in *pi6^em1/em1^* diplotene spermatocytes (*p* = 0.0001) and 1.5-fold in secondary spermatocytes (*p* = 0.007), and spermatids (*p* = 0.0005). A corresponding increase in piRNA precursor transcripts accompanied the decrease in piRNA abundance: in both *pi6^em1/em1^* and *pi6^em2/em2^* mutants, *pi10-qC2-545.1* precursors increased 2.1–3-fold in pachytene spermatocytes and diplotene spermatocytes (FDR = 3 × 10^−13^ to 6 × 10^−5^; Figure 7B and Table S2). Similarly, the abundance of *pi10-qA3-143.1* precursor transcripts increased 2–2.7-fold in diplotene spermatocytes, secondary spermatocytes, and spermatids (FDR = 4 × 10^−26^ to 2 × 10^−10^) in the two *pi6* mutant alleles.

**Figure 7.**
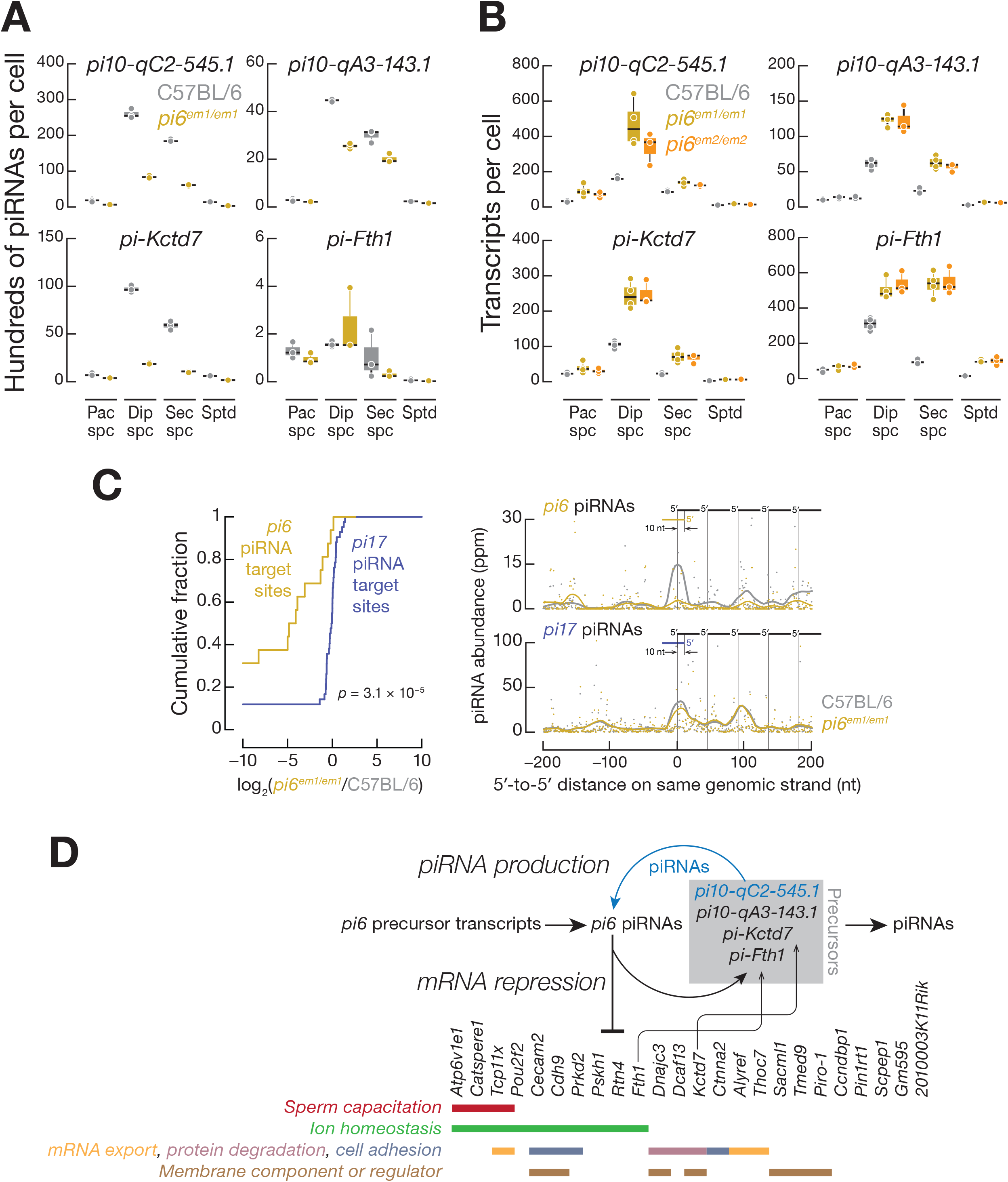
*pi6* piRNAs and piRNAs from other loci form a network to repress mRNA expression and facilitate piRNA biogenesis. **(A)** Abundance of mature piRNAs measured by small RNA-seq. Each dot represents the abundance of uniquely mapping reads in one biological sample. **(B)** Expression of piRNA precursors measured by RNA-seq. Each dot represents the abundance of transcripts in one biological sample. In **(A)** and **(B)**, horizontal black lines denote median; whiskers indicate 75^th^ and 25^th^ percentiles. **(C)** Analysis of 5′ to 5′ distances for mature piRNAs derived from pachytene piRNA precursors cleaved by *pi6* piRNAs. *pi6* piRNA-directed cleavage sites were identified requiring g2–g15 base-pairing between a *pi6* piRNA and a 5′ monophosphorylated, 3′ cleavage fragment (degradome-seq). *p* value was computed using the Kolmogorov-Smirnov test. **(D)** A model for *pi6* piRNA biogenesis and function. See also Figures S5 and S6, and Tables S4 and S5.

Further supporting the idea that *pi6* piRNA-directed cleavage initiates pachytene piRNA production from *pi10-qC2-545.1*, degradome sequencing detected two different *pi6* piRNA-dependent cleavage sites in *pi10-qC2-545.1* transcripts (Figure S5B; Table S4). Each cleavage site can be explained by an extensively complementary *pi6* piRNA predicted to direct MILI or MIWI to cut the *pi10-qC2-545.1* precursor transcript immediately before the 5′ end of the 5′ monophosphorylated RNA identified by degradome sequencing. For example, when cut by a *pi6* piRNA (10,604 molecules per cell) in wild-type diplotene spermatocytes, RNA fragments derived from a cleavage site at *chr10: 94,675,047–94,675,048* decreased >70-fold in *pi6^em1/em1^* diplotene spermatocytes (*n* = 2). For *pi10-qA3-143.1*, we identified a *pi6* piRNA-dependent cleavage site that produces 5′ monophosphorylated RNA in wild-type pachytene (3.3 ppm, *n* = 2) and diplotene spermatocytes (4.8 ppm, *n* = 2) but not in *pi6^em1/em1^* cells.

Consistent with *pi6* piRNAs initiating piRNA production by cleaving pachytene piRNA precursor transcripts, we detected a large reduction in the abundance of 3′ cleavage fragments from RNAs targeted by *pi6* piRNAs and mapping to pachytene piRNA loci in *pi6^em1/em1^* diplotene spermatocytes (16 cleavage sites), but not for RNAs targeted by *pi17* piRNAs (42 cleavage sites; Figure 7C, left panel). Strings of head-to-tail piRNAs (“phased” or “trailing” piRNAs) beginning at the 5′ end of a piRNA-directed 3′ cleavage fragment are the hallmark of piRNA-initiated piRNA production. We detected such phased piRNAs downstream of *pi6* piRNA-directed cleavage sites within pachytene piRNA precursors in wild-type diplotene spermatocytes (Figure 7C, right panel). In *pi6* mutants, the abundance of these trailing piRNAs decreased, whereas the abundance of trailing piRNAs initiated by *pi17* piRNAs was unchanged. Remarkably, *pi10-qC2-545.1* piRNAs whose production required *pi6* piRNAs reciprocally promote *pi6* piRNA biogenesis: three *pi10-qC2-545.1* piRNAs can account for *pi6* precursor cleavage sites that initiate *pi6* piRNA production in wild-type pachytene and diplotene spermatocytes (Figure S5C; Table S4). *pi10-qC2-545.1* is found in both mice and rats, but the gene produces a lncRNA in rats and a piRNA precursor in mice. The finding that *pi10-qC2-545.1* generates piRNAs only in *Mus musculus* suggests it emerged recently as a pachytene piRNA-producing locus. Perhaps the fortuitous production of *pi6* piRNAs with sufficient complementarity to direct cleavage of *pi10-qC2-545.1* transcripts has converted the locus to a source of piRNAs that enhance piRNA production from the more ancient *pi6* locus.

In addition to pachytene piRNA precursor targets, *pi6* piRNAs also initiate piRNA biogenesis from two piRNA-regulated protein-coding genes, *Kctd7* (*chr5: 130,144,861– 130,155,806* and *Fth1* (*chr19: 9,982,703–9,985,092*). These loci had been previously annotated as sources of hybrid (*pi-Kctd7*, chr5: 130,144,861–130,155,806) or pre-pachytene (*pi-Fth1*, chr19: 9,982,703–9,985,092) piRNAs. Like many piRNA-producing mRNAs, *Kctd7* produces piRNAs (9,700 ± 400 molecules per diplotene spermatocyte, *n* = 3) from its 3′ UTR. In contrast, *Fth1* piRNAs (160 ± 10 molecules per diplotene spermatocyte, *n* = 3) derive from the exons and 5′ UTR of the mRNA. In *pi6^em1/em1^* diplotene spermatocytes and secondary spermatocytes, the abundance of *Kctd7* piRNAs declined 5.3-fold (*p* < 0.0001, *n* = 3; Figure 7A), while the *Kctd7* and *Fth1* transcripts more than doubled in *pi6* mutants (Figure 7B; Table S2). Further supporting the idea that *pi6* piRNAs initiate piRNA biogenesis from these two protein-coding loci, our data identify two *pi6* piRNA-directed cleavage sites in the *Kctd7* 3′ UTR and two sites in exons (exon 2 and exon 4) of the *Fth1* mRNA (Table S4). Together, our data demonstrate that *pi6* piRNAs not only repress mRNA expression but also initiate piRNA biogenesis in *trans* from other piRNA-producing loci (Figure 7D).

## DISCUSSION

Deletion of the promoter of the mouse *pi6* pachytene piRNA locus causes specific, quantifiable defects in male fertility. These include impaired sperm capacitation and a failure of sperm to bind and penetrate the zona pellucida. The male fertility defects accompanying loss of *pi6* piRNAs are specific to this locus, as deletion of the promoter of *pi17*, which eliminates *pi17* piRNAs, had no detectable effect on male or female fertility or viability, as reported previously (Homolka et al., 2015). The phenotypic defects of *pi6* mutants reflect the molecular changes—increased steady-state abundance of mRNAs that encode proteins functioning in sperm capacitation, acrosome function, and other pathways with links to sperm biology. Pachytene piRNAs have been proposed to act collectively in meiotic spermatocytes or post-meiotic spermatids to target mRNAs for destruction (Gou et al., 2014; Goh et al., 2015).Our finding that deletion of *pi6*, but not of *pi17*, the most prolific piRNA-producing locus, leads to male fertility defects, suggests that individual pachytene piRNA loci can regulate distinct sets of genes. Our data argue against pachytene piRNAs acting en mass (Post et al., 2014), since not only does *pi6* produce far fewer piRNAs than *pi17*, but just a tiny fraction of *pi6* piRNAs can explain the effect of loss of *pi6* piRNAs on the transcriptome. *pi6* produces 80,354 distinct piRNA sequences, representing 10,943 unique g1–g21 piRNA sequences reproducibly detected (*n* = 3) present at >1 molecule per cell. Yet loss of *pi6* piRNAs dysregulates just 24 mRNAs, consistent with its remarkably specific mutant phenotype.

Moreover, our data argue strongly against miRNA-like regulation by piRNAs. If pachytene piRNAs found their target RNAs by a miRNA-like, seed-based mechanism, the predicted target repertoire of piRNAs produced by individual loci would be enormous: *pi6* piRNAs encompass 9,880 distinct 7mer-m8 seed sequences (g2–g8; Bartel, 2009), while *pi17* generates 134,358 distinct piRNA sequences, encompassing 11,324 distinct g2–g8 seeds. Yet loss of *pi17* piRNAs has no detectable phenotype, while loss of *pi6* piRNAs causes specific defects in sperm motility, the acrosome reaction, and egg fertilization. Although 104 *pi6* piRNAs are more abundant than miR-20a, the tenth most abundant miRNA in diplotene spermatocytes, loss of *pi6* piRNAs reproducibly increases the abundance of just 24 mRNAs. Of these, just six mRNAs appear to be the direct cleavage targets of *pi6* piRNAs, consistent with pachytene piRNAs acting like long siRNAs.

Finally, the phenotypic and molecular specificity of *pi6* mutants likely reflects a low degree of redundancy with other piRNA-producing loci. Nonetheless, other piRNA-producing loci may partially rescue loss of *pi6* piRNAs, accounting for the incomplete penetrance of the *pi6* sterility phenotype. Conversely, the lack of a phenotype for other pachytene piRNA-producing loci may simply reflect greater redundancy with their piRNA-producing peers. In this view, loss of regulation of the targets of *pi17* piRNAs may be compensated by piRNAs from other loci. Testing this hypothesis is clearly a prerequisite to explain why loss of *pi6* and not *pi17* piRNAs has an obvious biological consequence.

The current model for piRNA biogenesis in bilateral animals posits that piRNA-directed cleavage precursor transcripts facilitates the biogenesis of other piRNAs (Gainetdinov et al., 2018; Ozata et al., 2019). Consistent with this view, *pi6* piRNAs are required for biogenesis of piRNAs from four other piRNA-producing loci: *pi10-qC2-545.1* and *pi10-qC2-143.1*, both sources of pachytene piRNAs; and two protein-coding genes, *Kctd7*, a hybrid piRNA gene, and *Fth1*, a pre-pachytene piRNA gene. Despite producing just 3% as many piRNAs as *pi6*, *pi10-qC2-545.1* makes piRNAs that can cleave *pi6* transcripts, initiating biogenesis of *pi6* piRNAs. It is tempting to speculate that such positive feedback loops operate among many piRNA-producing loci. The extreme sequence diversity of piRNAs may serve mainly to initiate robust piRNA production among the pachytene piRNA loci rather than to regulate the abundance of large numbers of mRNAs.

Beyond the requirement for *pi6* piRNAs to produce fully functional sperm, *pi6* piRNAs appear to play an additional role in embryo development. Our data suggest that the arrested development and reduced viability of embryos derived from *pi6* mutant sperm reflects a paternal defect and not the embryonic genotype. Damaged sperm DNA, abnormal sperm chromatin structure, and failure to form a male pronucleus in fertilized embryos have been reported to be linked to retarded embryo development (Sakkas et al., 1998; Borini et al., 2006). Our analysis of transposon RNA abundance in *pi6* mutant germ cells argues against a role for *pi6* piRNAs in transposon silencing during spermatogenesis, but we cannot currently exclude a direct or indirect role for *pi6* piRNAs in silencing transposons in the early embryo (Peaston et al., 2004). Of course, DNA damage might reflect incomplete repair of the double-stranded DNA breaks required for recombination, rather than transposition or transposon-induced illegitimate recombination.

How pachytene piRNAs identify their targets remains poorly understood, in part because of a lack of suitable biochemical or genetic model systems. The availability of a mouse mutant missing a specific set of piRNAs whose absence causes a readily detectable phenotype should provide an additional tool for understanding the base-pairing rules that govern the binding of piRNAs to their RNA targets and for unraveling the regulatory network created by pachytene piRNAs.

## SUPPLEMENTAL INFORMATION

Supplemental Information includes Extended Experimental Procedures, Figures S1–6, Tables S1–S5, and Movies S1–S10.

## AUTHOR CONTRIBUTIONS

P.-H.W., K.C., Y.F., Z.W., and P.D.Z. conceived and designed the experiments. P.-H.W., K.C., D.M.Ö., A.A., and C.C. performed the experiments. Y.F., T.Y., I.G., and P.-H.W. analyzed the sequencing data. P.-H.W., Y.F., and P.D.Z. wrote the manuscript.

## ACKNOWLEDGEMENTS

We thank P. Cohen, K. Grive, and E. Crate at Cornell University for generously sharing protocols and advice on germ cell sorting and meiotic chromosome studies; H. Florman, P. Visconti, and M. Gervasi for sharing protocols and advice on sperm studies; the UMMS Transgenic Animal Modeling Core for advice on fertility test and embryo phenotype; the UMMS FACS core for advice on and help with germ cell sorting; the UMMS EM Core (supported by National Center for Research Resources Award SI0OD021580) for advice on and help with sperm transmission electron microscopy; and members of our laboratories for critical comments on the manuscript. This work was supported in part by National Institutes of Health grants GM65236 to P.D.Z. and P01HD078253 to P.D.Z. and Z.W.

## STAR METHODS

### Mouse mutants

Mice were maintained and sacrificed according to guidelines approved by the Institutional Animal Care and Use Committee of the University of Massachusetts Medical School (A-2222-17).

Small guide RNAs (sgRNAs) flanking piRNA promoters were designed using CRISPR design tools (crispr.mit.edu/). DNA oligos containing guide sequences were cloned into pX330 vectors (Cong et al., 2013), and their cleavage activity tested in NIH3T3 cells by co-transfecting pX330 constructs containing sgRNA sequences and puromycin-resistant plasmid (pPUR) using TransIT-X2 (Mirus Bio, Madison, WI). Puromycin (3 µg/µl) was added 24 h after transfection and DNA extracted 48 h afterwards. Promoter deletions were detected by PCR using primers flanking the predicted Cas9 cleavage sites.

For mice, sgRNAs were generated by in vitro transcription and purified by electrophoresis on 8% (w/v) polyacrylamide gels. To generate the *pi6^em1/em1^* and *pi17^−/−^* lines used in this study, in vitro transcribed sgRNAs (10 ng/µl each) targeting *pi6* and *pi17* were mixed with Cas9 mRNA (40 ng/µl) and injected together into the cytoplasm of one-cell C57BL/6 zygotes (RNA only). For some founders, the sgRNA and Cas9 mRNA mixture was combined with pX330 plasmids expressing the same four sgRNAs and Cas9 and injected into both the cytoplasm and pronuclei of one-cell C57BL/6 zygotes (RNA + DNA). For *pi6^em2/em2^*, in vitro transcribed sgRNAs and Cas9 mRNA were injected into the cytoplasm of one-cell C57BL/6 embryos. Embryos were transferred to pseudopregnant females using standard methods. To screen for mutant founders, DNA was extracted from small pieces of tail clipped from three-week-old pups (Truett et al., 2000). Deletions were detected by PCR, and PCR products purified and cloned into TOPO blunt vectors. Mutant sequences were determined by Sanger sequencing. Mouse mutant lines were established and maintained by mating mutant founders with C57BL/6 males or females. All mutant mice in this study were backcrossed for at least two generations before use.

### Mouse fertility test

Each 2–8 month-old male mouse was housed with one 2–4 month-old C57BL/6 female, who was examined for the presence of a vaginal plug the following morning. When a plug was observed, the female was housed separately. For male mice who did not produce pups after 3 months (∼3 cycles), the original female was replaced with a new female and the fertility test continued.

### Testis histology, sperm count, and sperm morphology

Mouse testes were fixed in Bouin’s solution overnight, washed with 70% ethanol, embedded in paraffin, and sectioned at 5 µm thickness. Sections were stained with hematoxylin solution, countered stained with eosin solution, and imaged using Leica DMi8 brightfield microscope equipped with an 20× 0.4 N.A. objective (HC PL FL L 20×/0.40 CORR PH1, Leica Microbiosystems, Buffalo Grove, IL). To quantify sperm abundance, the cauda epididymides were collected from mice and placed in phosphate-buffered saline (PBS) containing 4% (w/v) bovine serum albumin. A few incisions were made in the epididymides with scissors to release the sperm, followed by incubation at 37°C and 5% CO_2_ for 20 min. A 20 µl aliquot of sperm suspension was diluted in 480 µl of 1% (w/v) paraformaldehyde (PFA), and sperm cells counted at 10× by brightfield microscopy. To assess sperm morphology, caudal epididymal sperm were fixed in 1% (w/v) PFA, stained with trypan blue, and a Leica DMi8 brightfield microscope equipped with an 63× 1.4 N.A. oil immersion objective (HC PL APO; Leica Microbiosystems, Buffalo Grove, IL). Sperm stained with Alexa 488-conjugated PNA (see below) were also used to assess sperm morphology.

### Meiotic chromosome spreads

Meiotic chromosome spreads were prepared as described (Holloway et al., 2014). Mouse testes were incubated in hypotonic buffer (30 mM Tris-Cl, pH 8.2, 50 mM sucrose, 17 mM sodium citrate, 5 mM EDTA, 0.5 mM DTT) for 30 min on ice, then small fragments of seminiferous tubules were moved to 100 mM sucrose solution and pulled apart with forceps to release germ cells. A drop of sucrose solution containing germ cells was pipetted onto a glass slide with a thin layer of 1× PBS containing 1% PFA and 0.15% (v/v) Triton-X100 (pH 9.2) and spread by swirling. Slides were placed in a humidifying chamber for 2.5 h, air-dried, and washed twice with 1× PBS with 0.4% Photo-Flo 200 (Kodak, Rochester, NY) and once with water with 0.4% Photo-Flo 200, and air-dried. For immunostaining of meiotic chromosomes, slides were sequentially washed with (1) 1× PBS with 0.4% Photo-Flo 200, (2) 1× PBS containing 0.1% (v/v) Triton-X, and (3) blocked with PBS containing 3% (w/v) BSA, 0.05% (v/v) Triton X-100, and 10% (v/v) goat serum in 1× PBS at room temperature. The slides were then incubated with primary antibodies, anti-SCP1 (1:1000 dilution) and anti-SCP3 (1:1000 dilution), in a humidifying chamber overnight at room temperature. Washing and blocking steps were repeated the next day, and the slides were incubated with Alexa 488-or Alexa 594-conjugated secondary antibodies (1:10,000 dilution) for 1 h at room temperature. Slides were washed thrice with 1× PBS containing 0.4% (v/v) Photo-Flo 200, once with water containing 0.4% Photo-Flo 200 mixture, air-dried in the dark, mounted by incubation in ProLong Gold Antifade Mountant with DAPI (4ʹ,6ʹ-diamidino-2-phenylindole; Thermo Fisher Scientific, Waltham, MA) overnight in the dark, and imaged using a Leica DMi8 fluorescence microscope equipped with an 63× 1.4 N.A. oil immersion objective (HC PL APO; Leica Microbiosystems, Buffalo Grove, IL).

### Cell sorting by FACS

Testicular cell sorting was performed as described (Cole et al., 2014). Testes were collected, decapsulated, and incubated in 0.4 mg/ml collagenase type IV (Worthington LS004188) in 1× Grey′s Balanced Salt Solution (GBSS, Sigma, G9779) at 33°C rotating at 150 rpm for 15 min. Separated seminiferous tubules were washed with 1× GBSS and incubated in 0.5 mg/ml Trypsin and 1 µg/ml DNase I in 1× GBSS at 33°C rotated at 150 rpm for 15 min. Tubules were dissociated on ice by gentle pipetting, and then 7.5% (v/v) fetal bovine serum (f.c.) was added to inactivate trypsin. The cell suspension was filtered through a pre-wetted 70 µm cell strainer, and cells pelleted at 300 × *g* for 10 min at 4°C. Cells were resuspended in 1× GBSS containing 5% (v/v) FBS, 1 µg/ml DNase I, and 5 μg/ml Hoechst 33342 (Thermo Fisher Scientific, Waltham, MA) and rotated at 150 rpm at 33°C for 45 min. Propidium iodide (0.2 μg/ml, f.c.; Thermo Fisher Scientific, Waltham, MA) was added, and cells strained through a pre-wetted 40 µm cell strainer. Cell sorting was performed on a FACSAria II (BD Biosciences, Franklin Lakes, NJ). The purity of sorted fractions was assessed by immunostaining. Secondary spermatocyte and spermatid populations were >90% pure, and the pachytene spermatocytes and diplotene spermatocytes were >80% pure.

### In vitro fertilization (IVF) and embryo transfer

In vitro fertilization was performed as previously described (Nagy et al., 2003) using spermatozoa from caudal epididymis of C57BL/6, *pi6^+/em1^*, or *pi6^em1/em1^* mice. Spermatozoa were incubated in complete human tubal fluid media (HTF; 101.6 mM NaCl, 4.69 mM KCl, 0.37mM KH_2_PO_4_, 0.2 mM MgSO_4_⋅7H_2_O, 21.4 mM Na-lactate, 0.33 mM Na-pyruvate, 2.78 mM glucose, 25 mM NaHCO_3_, 2.04 mM CaCl_2_⋅2H_2_O, 0.075 mg/ml Penicillin-G, 0.05 mg/ml streptomycin sulfate, 0.02% (v/v) phenol red, 4 mg/ml BSA) with oocytes (98–146 for control sperm and 120–293 for *pi6^em1/em1^* sperm) from B6SJLF1/J mice for 3–4 h at 37°C with constant 5% O_2_, 90% N_2_, and 5% CO_2_ concentration. Oocyte viability and the presence of pronuclei were assessed under a Nikon SMZ-2B (Nikon, Tokyo, Japan) dissecting microscope. To observe embryo development, embryos were moved into potassium-supplemented simplex optimized media (KSOM; 95 mM NaCl, 2.5 mM KCl, 0.35 mM KH_2_PO_4_, 0.2 mM MgSO_4_⋅7H_2_O, 10 mM Na-lactate, 0.2 mM Na-pyruvate, 0.2 mM glucose, 25 mM NaHCO_3_, 1.71 mM CaCl_2_⋅2H_2_O, 1 mM L-glutamine, 0.01 mM EDTA, 0.075 mg/ml Penicillin-G, 0.05 mg/ml streptomycin sulfate, 0.02% (v/v) phenol red, 1 mg/ml BSA; Millipore Sigma, Burlington, MA) after IVF and assessed every 24 h. To measure birth rates, two-cell embryos were transferred to Swiss Webster pseudopregnant females, and fetuses isolated by cesarean section 18.5 d after embryo transfer.

For zona-free IVF, the zona pellucida of oocytes was removed with acid Tyrode’s solution as described (Yanagimachi et al., 1976; Johnson et al., 1991).

### Intracytoplasmic sperm injection (ICSI)

Frozen caudal epididymal spermatozoa were thawed, the sperm tails detached (Nagy et al., 2003), and individual *pi6^+/em1^* or *pi6^em1/em1^* sperm heads injected into B6D2F1/J oocytes in Chatot-Ziomek-Bavister media (CZB; 81.62 mM NaCl, 4.83 mM KCl, 1.18 mM KH_2_PO_4_, 1.18 mM MgSO_4_⋅7H_2_O, 25 mM Na_2_HCO_3_, 1.70 mM CaCl_2_⋅2H_2_O, 0.11 mM Na_2_-ETDA⋅2H_2_O, 1 mM L-glutamine, 28 mM Na-lactate, 0.27 mM Na-pyruvate, 5.55 mM glucose, Penicillin-G 0.05 mg/ml, 0.07 mg/ml streptomycin sulfate, 4 mg/ml BSA; Millipore Sigma, Burlington, MA) using the PiezoXpert (Eppendorf, Hamburg, Germany; Cat#5194000024). Surviving oocytes were counted, collected, and cultured in KSOM (Millipore Sigma, Burlington, MA) at 37°C and 5% CO_2_ for 24 h. Two-cell embryos were surgically transferred unilaterally into the oviducts of pseudopregnant Swiss Webster females. At 16.5 days after the surgery, live fetus isolated by cesarean section.

### Sperm motility

Cauda epidydimal sperm were collected from mice and placed in 37°C HTF media containing 4 mg/ml BSA in an incubator with 5% CO_2_. A drop of sperm was removed from the suspension and pipetted into a sperm counting glass chamber, then assayed by CASA or video acquisition. CASA was conducted using an IVOS II instrument (Hamilton Thorne, Beverly, MA) with the following settings: 100 frames acquired at 60 Hz; minimal contrast = 50; 4 pixel minimal cell size; minimal static contrast = 5; 0%straightness (STR) threshold; 10 μm/s VAP Cutoff; prog. min VAP, 20 μm/s; 10 μm/s VSL Cutoff; 5 pixel cell size; cell intensity = 90; static head size = 0.30–2.69; static head intensity = 0.10–1.75; static elongation = 10–94; slow cells motile = yes; 0.68 magnification; LED illumination intensity = 3000; IDENT illumination intensity = 3603; 37°C. The raw data files (i.e. .dbt files for motile sperm and .dbx files for static sperm) were used for sperm motility analysis. For the motile sperm, only those whose movement was captured with ≥45 consecutive frames were analyzed. For the boxplots, the number of static sperm was re-calculated for each mouse according to the percentage of motile sperm with ≥ 45 frames. For hyperactivated motility analysis, .dbt files of motile sperm were used as input for CASAnova, as previously described (Goodson et al., 2011). For movie acquisition, a Nikon Diaphot 200 microscope (Nikon, Tokyo, Japan) with darkfield optics equipped with Nikon E Plan 10×/0.25 160/-Ph1 DL objective (Nikon, Tokyo, Japan), ZWO ASI 174mm Monochrome CMOS Imaging camera (ZWO, SuZhou, China), and the SharpCap software (https://docs.sharpcap.co.uk/2.9/) using darkfield at 10× magnification were used to record sperm movement at 37°C.

### In vitro acrosome reaction and capacitation assay

Sperm capacitation was induced and acrosome reaction was assessed as described (Talbot et al., 1976). Cauda epididymides were collected from mice, placed in HTF media containing 4 mg/ml BSA pre-warmed for at least 2 h in a 37°C incubator at 5% CO_2_. A few incisions were made in the epididymides with scissors to release the sperm, followed by incubation at 37°C in 5% CO_2_ for 90 min. Calcium ionophore A23187 (10 µm f.c. in DMSO) was added, and incubation continued for 30 min. Sperm were fixed at room temperature for 10 min by adding two volumes of 4% (w/v) PFA, pelleting at 1,000 × *g* for 5 min, washed with 1× PBS, resuspended in fresh 1× PBS, spotted on a glass slide, and air-dried. Methanol was pipetted onto the sperm to permeabilize the cells, followed by washing with 1× PBS. Slides were incubated overnight in 10 µg/ml Alexa Fluor 488-conjugated peanut agglutinin (PNA) in 1× PBS (Mortimer D., 1987), washed with 1× PBS, air-dried, and mounted with ProLong Gold Antifade Mountant with DAPI (Thermo Fisher Scientific, Waltham, MA). Sperm were imaged using a Leica DMi8 fluorescence microscope equipped with a 63× 1.4 N.A. oil immersion objective (HC PL APO; Leica Microbiosystems, Buffalo Grove, IL) and analyzed using ImageJ (version 2.0.0-rc-68/1.52e; https://fiji.sc/).

### Transmission electron microscopy

Mouse caudal epididymides were dissected and immediately fixed by immersion in Karnovsky’s fixative (2% formaldehyde (v/v) and 3% glutaraldehyde (v/v) in 0.1M sodium phosphate buffer, pH 7.4; Electron Microscopy Sciences, Hatfield, PA) overnight at 4°C, and washed three times in 0.1M phosphate buffer. Following the third wash, the tissues were post-fixed in 1% osmium tetroxide (w/v; Electron Microscopy Sciences, Hatfield, PA) for 1 h at room temperature, washed three more times with water for 10 min each, and dehydrated using a graded series of 30%, 50%, 70%, 85%, 95%, 100% (3 changes) ethanol and 100% propylene oxide (two changes) and a mixture of 50% propylene oxide (v/v) and 50% SPI-Pon 812 resin mixture (v/v; SPI Supplies, West Chester, PA). The sample was incubated in seven successive changes of SPI-Pon 812 resin over three days, polymerized at 68°C in flat molds, and reoriented to allow cross-sectioning of spermatozoa in the lumen of epididymis. 70nm sections were cut on a Leica EM UC7 ultramicrotome (Leica Microsystems, Wetzlar, Germany) using a diamond knife, collected on copper mesh grids, and stained with 3% lead citrate (w/v) and 0.1% uranyl acetate (w/v) to increase contrast. Finally, sections were examined using Philips CM10 transmission electron microscope (Philips Electron Optics, Eindhoven, The Netherlands) at 100 KV. Images were recorded using the Erlangshen digital camera system (Gatan Inc., Pleasanton, CA).

### RNA-seq and small RNA-seq analysis

Small RNA-seq and RNA-seq libraries were constructed incorporating unique molecular identifiers for removal of PCR duplicates and sequenced using NextSeq 500 (Illumina, San Diego, CA) as described (Fu et al., 2018; Gainetdinov et al., 2018). To sequence mature piRNAs, small RNA was oxidized with 25 mM NaIO_4_ in 30 mM sodium borate, 30 mM boric acid (pH 8.6; Sigma Aldrich, St. Louis, MO) at 25°C for 30 min. RNA was precipitated with ethanol before adapter ligation. A set of 9 synthetic 2′-*O*-methylated RNA oligonucleotides was added to each RNA sample to allow measurement of molecules per cell. Small RNA-seq and RNA-seq reads were mapped to mouse genome assembly mm10 using piPipes (Han et al., 2015b). For small RNA quantification, sequences of synthetic spike-in oligonucleotides were identified allowing no mismatches and the number of molecules of small RNAs per library was calculated based on the read abundance of the spike-in oligonucleotides. For long transcript quantification, 1 µL of 1:100 dilution of ERCC spike-in mix 1 (Thermo Fisher, 4456740, LOT00418382) was added to 1 µg of total RNA in the first step. Differentially expressed transcripts were determined using DESeq2 (Love et al., 2014). Transcript abundance between *pi6^+/em1^* and C57BL/6 testes were indistinguishable (<2-fold change and FDR >0.05).

### Analysis of piRNA cleavage sites

Cleaved RNA fragments bearing 5′ monophosphates (“degradome” sequences) were cloned as previously described (Addo-Quaye et al., 2008; Wang et al., 2015) and sequenced using a paired-end sequencing kit on NextSeq 500 (Illumina, San Diego, CA). Briefly, 5′ monophosphate-bearing RNAs were enriched using an adapter with a 3′ hydroxyl and T4 RNA ligase. cDNA was generated using random primers conjugated with the sequence of the 3′ adapter and PCR-amplified using primers containing multiplex barcodes. piRNAs (>1 ppm) and degradome sequences from the same cell types were used for piRNA target analysis. Degradome sequences were extended 3′ to 5′ based on the mouse reference genome mm10, and putative targeting piRNAs were identified first by their pairing with specific degradome sequences at g2–g7 and a minimum of 8 additional base-pairs 5′ of g7. Only degradome sequences that begin at g11 of the matching piRNAs were used. Lastly, cleavage sites for which the read abundance significantly decreased (>2-fold and FDR <0.05) in *pi6* mutants were extracted. Analysis of 5′ to 5′ distance for mature *pi6* piRNAs was performed as described (Gainetdinov et al., 2018). Briefly, 5′ monophosphorylated pachytene piRNA precursor fragments from wild-type diplotene spermatocyte were detected by degradome-seq. Those fragments whose 5′ ends could be by explained by cleavage directed by complementary (g2—g15) *pi6* piRNAs were identified. Only fragments whose abundance decreased in *pi6* mutant diplotene spermatocytes were retained.

### qRT-PCR RNA quantification

Isolated total RNA from sorted germ cells was treated with Turbo DNase (Thermo Fisher Scientific, Waltham, MA) at 37°C for 30 min and purified using RNA Clean & Concentrator (Zymo Research, Irvine, CA). First strand cDNA was synthesized using oligo dT_20_ (for full-length transcripts) or random hexamers (for all transcript fragments) and Superscript III (Invitrogen, Carlsbad, CA). Quantitative PCR was performed for each sample using SsoFast EvaGreen Supermix (Bio-Rad, Hercules, CA) with technical triplicates. Relative transcript abundance was calculated using the ΔΔCt method. *Gapdhs*, a testis-specific mRNA that remains unchanged in *pi6* mutants based on RNA-seq analysis, was used for normalization. Statistical significance was calculated using the unpaired t-test.

### Transposon mapping

RNA-seq reads were intersected using BEDtools (Quinlan and Hall, 2010) with Repeat Masker annotation from UCSC (downloaded from https://genome.ucsc.edu/cgi-bin/hgTables). Reads mapping to multiple genomic locations were apportioned. Reads for individual repeats were aggregated to obtain reads counts for repeat families.

### Statistics

All statistics were performed using R (https://www.rstudio.com/) and graphs were generated using Igor Pro v7.08 (WaveMetrics) or ggplot2 v3.0.0 (https://ggplot2.tidyverse.org/). Unless otherwise stated, Mann-Whitney-Wilcoxon test was used to calculate *p* values.

## ACCESSION NUMBERS

All sequencing data are available through the NCBI Sequence Read Archive using accession number PRJNA480354.

## SUPPLEMENTAL FIGURE, TABLE, AND MOVIES

### Supplemental Figure Legends

**Figure S1.**
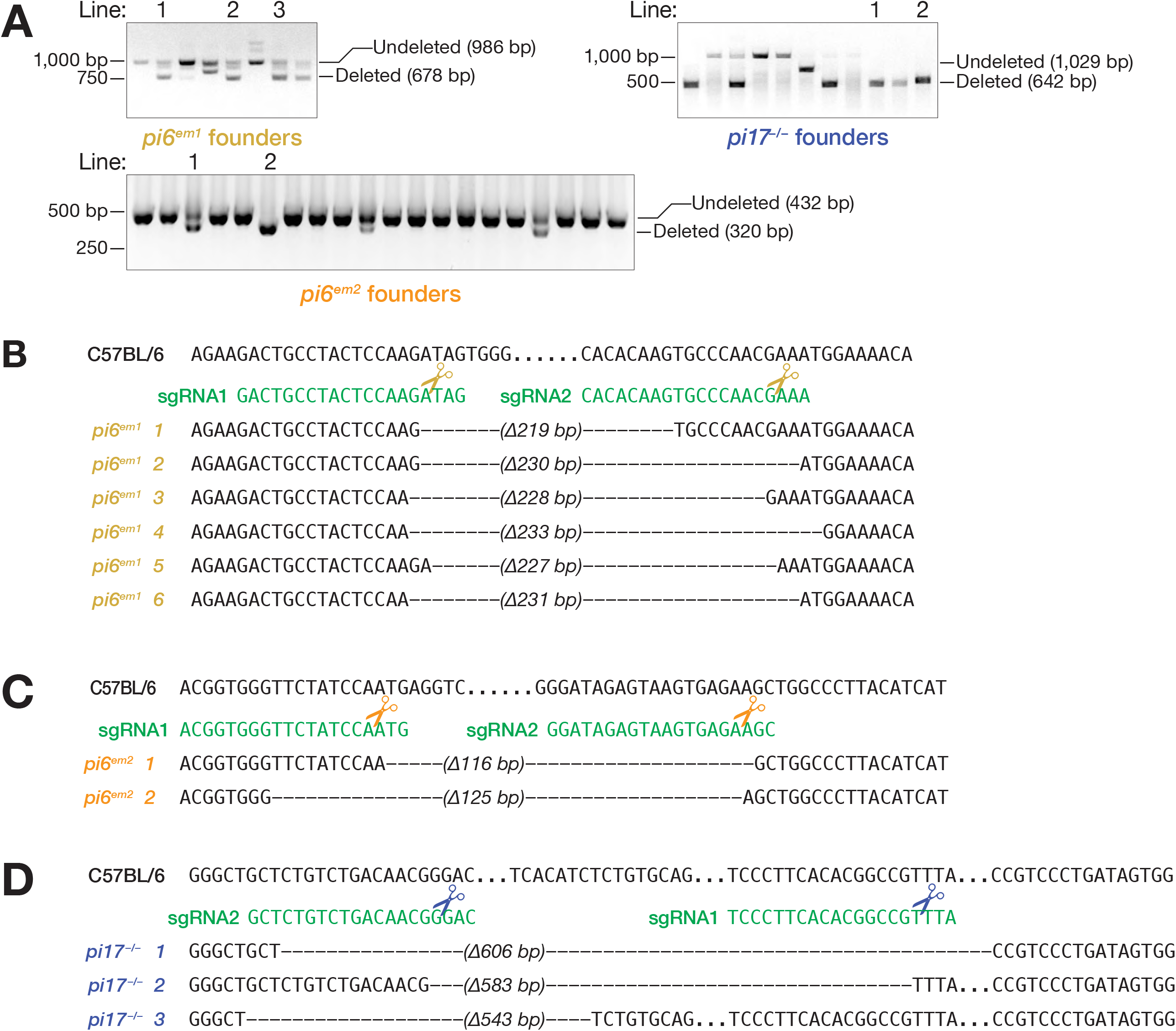
Confirmation of mutant founder genotypes. Related to Figure 1 and Table S1. **(A)** Genotyping of mutant founders by PCR. Genomic sequences of *pi6* promoter region in *pi6^em1^* **(B)** and *pi6^em2^* **(C)** mouse lines. **(D)** Genomic sequences of *pi17* promoter region in *pi17^−/−^* mouse lines. Dashes, genomic sequences deleted by CRISPR; dots, unaltered sequence omitted for clarity.

**Figure S2.**
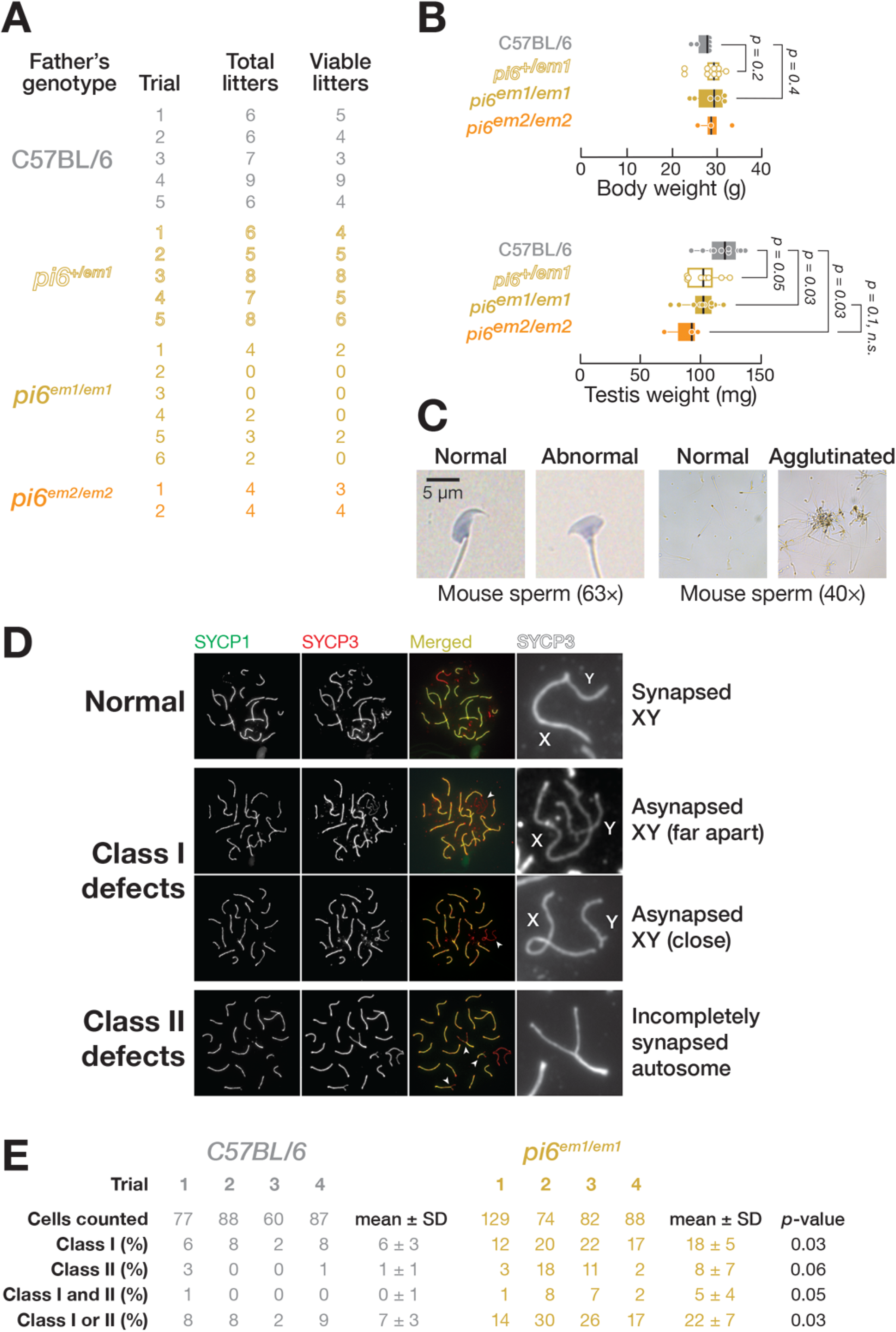
*pi6^em1/em1^* adult male phenotype. Related to Figure 2. **(A)** Number of litters produced in 6 months by 2–8 month-old males. **(B)** Body and testis weight of 2–4 month-old *pi6^em1/em1^* and *pi6^em2/em2^* males. Each dot represents an individual mouse. The thick lines denote median values, and whiskers indicate the 75^th^ and 25^th^ percentiles. **(C)** Representative spermatozoa. **(D)** Representative patterns of meiotic chromosome synapsis in *pi6^em1/em1^* pachytene spermatocytes. SYCP1, Synaptonemal complex protein 1; SYCP3, Synaptonemal complex protein 3. **(E)** Quantification of patterns of meiotic chromosome synapsis depicted in **(D)**.

**Figure S3.**
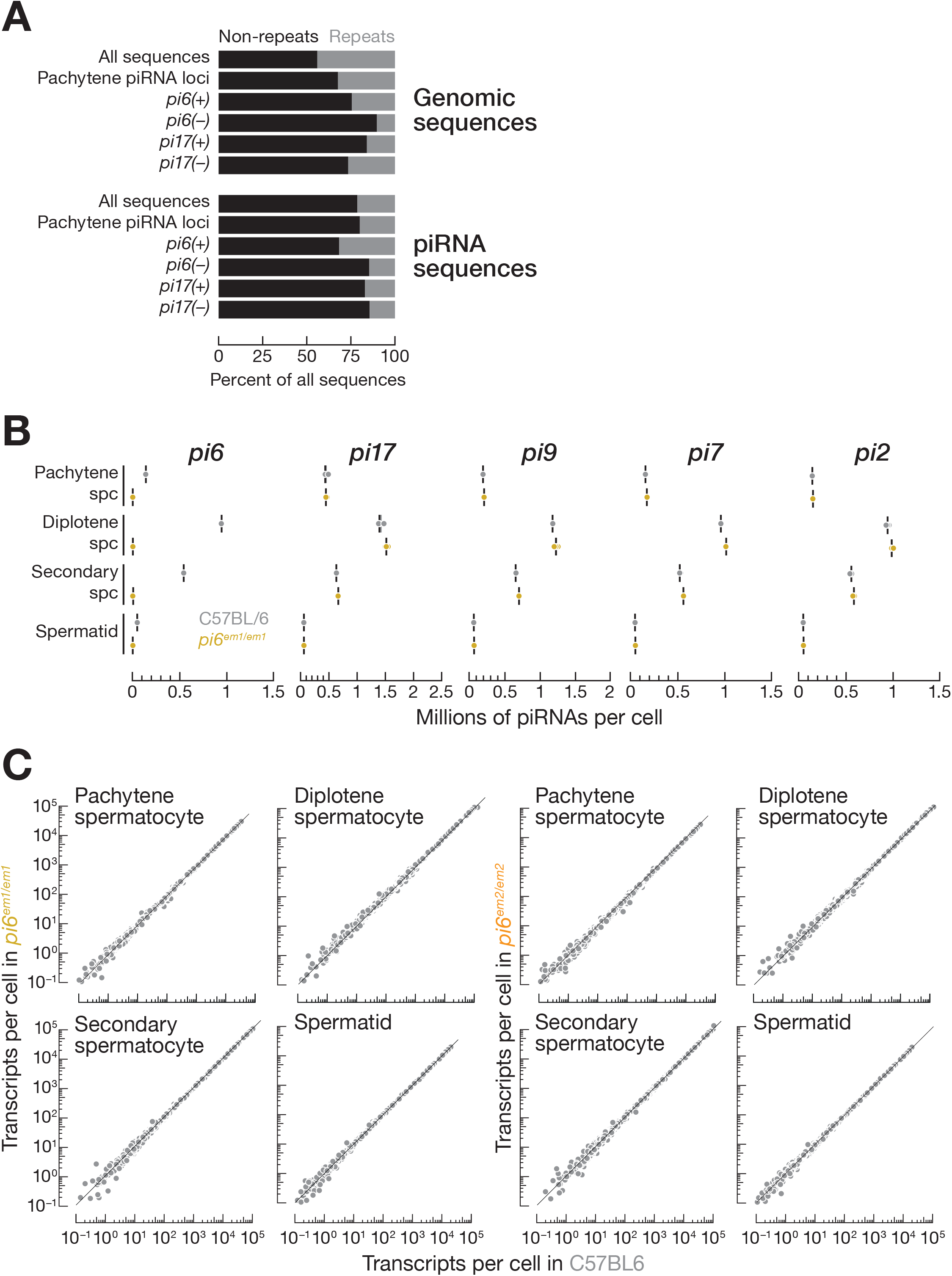
Abundance of transposons in *pi6^em1/em1^* and *pi6^em2/em2^* germ cells. Related to Figure 3. **(A)** Proportions of the whole genome or piRNA sequences composed of repetitive sequences. **(B)** Abundance of mature piRNAs from the top five major pachytene piRNA-producing loci in indicated cell types measured by small RNA-seq. Each dot represents the abundance of unique-mapping reads in one biological sample. Vertical black lines denote median; whiskers indicate 75^th^ and 25^th^ percentiles. **(C)** Abundance of transposon-derived RNAs in mouse germ cells. Each dot represents the mean of four (wild-type and *pi6^em1/em1^*) or three (*pi6^em2/em2^*) biologically independent RNA-seq experiments. Gray dots indicate change in abundance <2-fold and/or FDR > 0.05.

**Figure S4.**
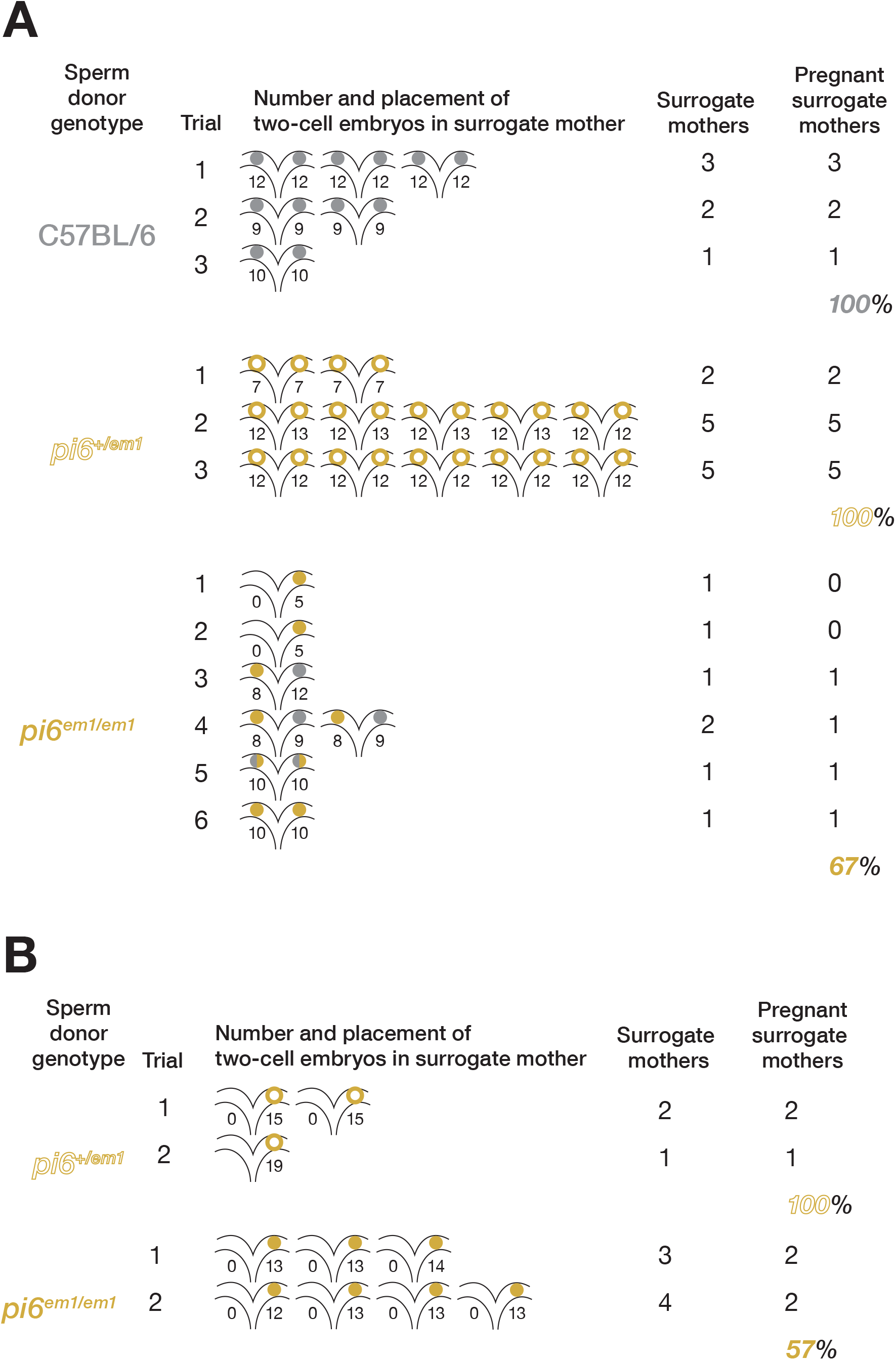
Pregnancy rate of surrogate mothers in IVF and ICSI experiments. Related to Figure 5. Percent of pregnant surrogate mothers in IVF **(A)** and ICSI **(B)**.

**Figure S5.**
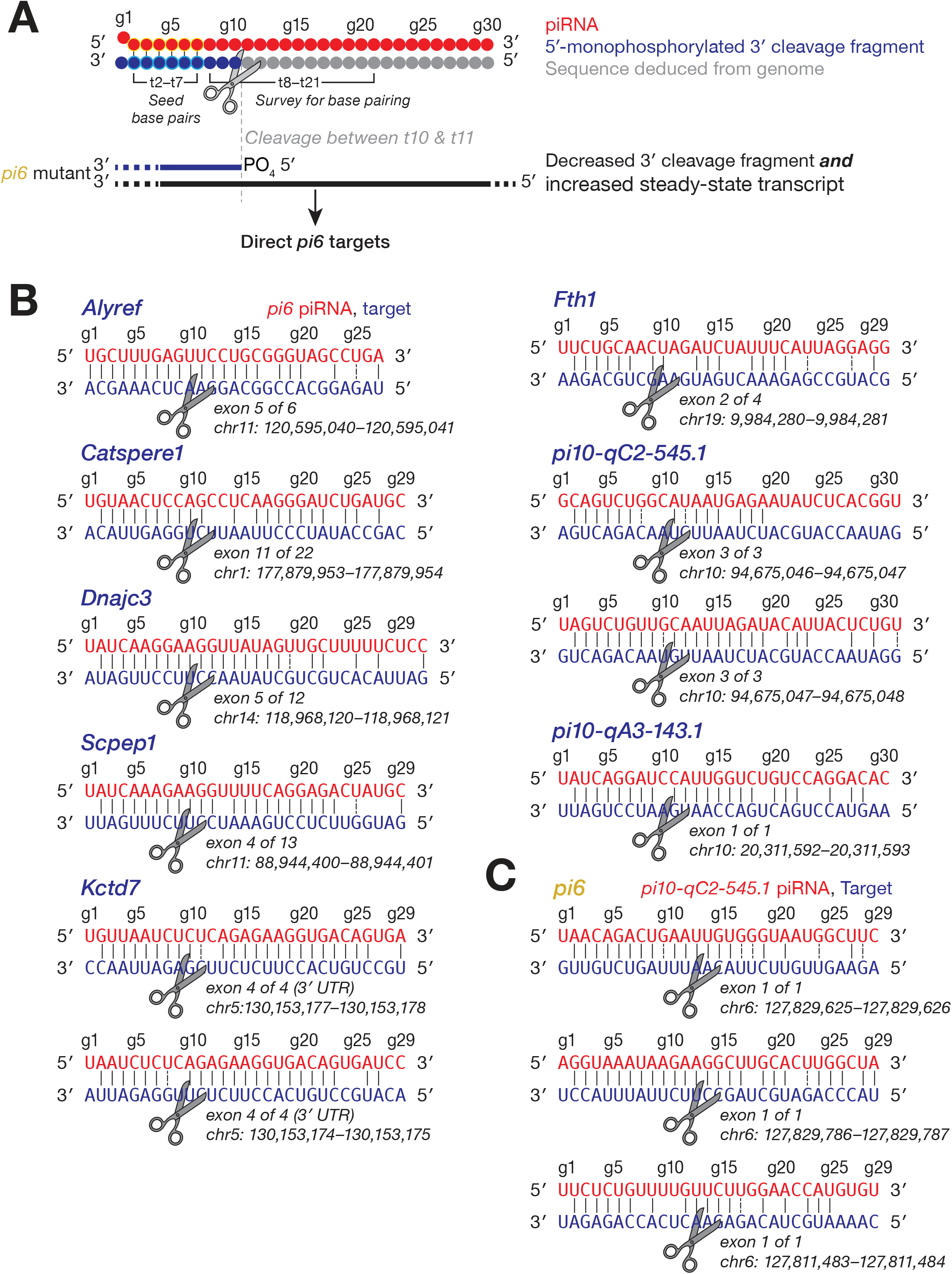
Transcripts directly cleaved by *pi6* and *pi10-qC2-545.1* piRNAs. Related to Figure 6 and Figure 7. **(A)** Strategy to identify piRNA-directed cleavage sites. **(B)** *pi6*-dependent cleavage sites in mRNAs or pachytene piRNA precursors from *pi10-qC2-545.1* and *pi10-qA3-143.1* showing inferred base pairing with the corresponding *pi6* piRNA guides. An exemplary piRNA guide is shown where more than one piRNA can direct the same cleavage. **(C)** Cleavage sites in *pi6* precursors explained by *pi10-qC2-545.1* piRNAs. An exemplary piRNA guide is shown.

**Figure S6.**
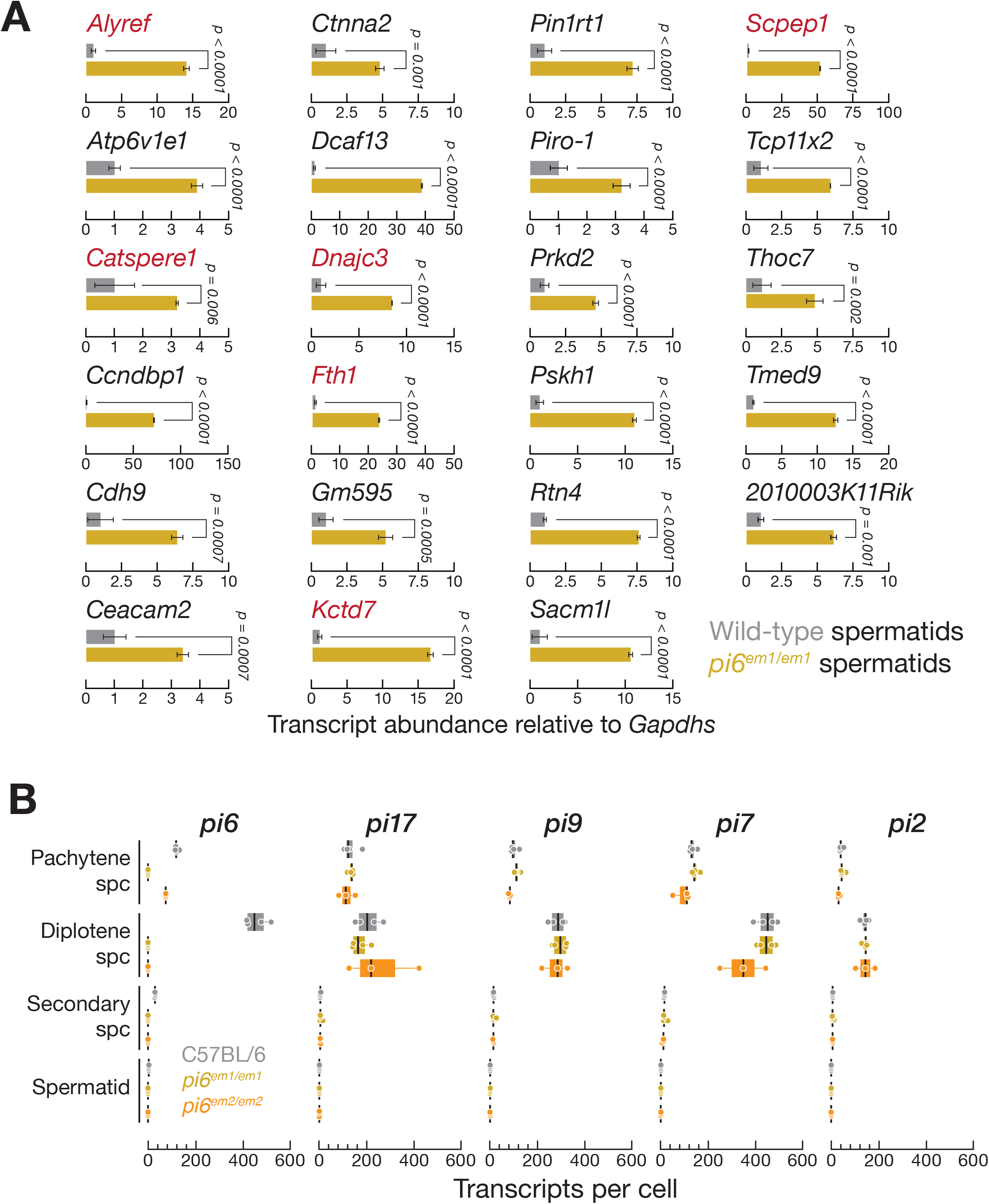
Transcriptome changes in *pi6^em1/em1^* and *pi6^em2/em2^* cells. Related to Figure 6 and Figure 7. **(A)** Expression of mRNAs measured by qRT-PCR using oligo dT_(20)_ to prime cDNA synthesis and PCR primers spanning *pi6* piRNA-directed cleavage sites (gene names in red) or designed to detect full-length RNA (gene names in black). *Pou2f2* mRNA abundance in spermatids was below the limit of detection by qRT-PCR. Vertical black line: mean ± S.D. (*n* = 3). **(B)** Abundance of piRNA precursors from the top five major pachytene piRNA-producing loci in indicated cell types measured by RNA-seq. Each dot represents the abundance of transcripts in one biological sample. Thick lines denote median; whiskers indicate 75^th^ and 25^th^ percentiles.

### Supplemental Table Legends

**Table S1.** Statistics for CRISPR/Cas9 genome editing to generate *pi6* mutants. Related to Figure 1 and S1.

| Allele | *pi6<sup>em1</sup>* |  |  |  | *pi6<sup>em2</sup>* |
| --- | --- | --- | --- | --- | --- |
| Nucleic acid injected | sgRNA + Cas9 mRNA | pX330 construct | sgRNA + Cas9 mRNA + pX330 construct | Total | sgRNA + Cas9 mRNA |
| Number of pups screened | 55 | 45 | 42 | 142 | 23 |
| Number of founders | 5 (9%) | 1 (2%) | 2 (5%) | 8 (6%) | 5 (22%) |
| Number of female founders | 3 (60%) | 1 (100%) | 1 (50%) | 5 (63%) | 2 (40%) |
| Number of male founders | 2 (40%) | 0 (0%) | 1 (50%) | 3 (38%) | 3 (60%) |
| Number of surviving founders | 5 (100%) | 0 (0%) | 2 (100%) | 7 (88%) | 5 (100%) |

**Table S2. Differentially expressed genes in *pi6^em1/em1^* and *pi6^em2/em2^* germ cells. Related to Figure 6 and S5.** Mean abundance (molecules per cell) of significantly altered mRNAs (>2-fold change ∩ FDR <0.05) in C57BL/6 (*n* = 4) versus *pi6^em1/em1^* (*n* = 4) and *pi6^em2/em2^* (*n* = 3) cells for RNA-seq replicate datasets. DESeq2 was used to calculate changes in transcript abundance and FDR.

**Table S3. Reported loss-of-function phenotype of mRNAs whose abundance increased in *pi6^em1/em1^* and *pi6^em2/em2^* germ cells. Related to Figure 6 and S2**. Reported mouse mutant phenotype or associated human fertility defects were obtained from Mouse Genome Informatics (http://www.informatics.jax.org/), International Mouse Phenotyping Consortium (https://www.mousephenotype.org/), or cited papers.

**Table S4. piRNA target sites identified using RNA-, small RNA-, and degradome-seq. Related to Figure 6 and 7** Direct piRNA targets identified using sequences of piRNAs and 5′ monophosphorylated 3′ cleavage RNA fragments.

**Table S5.**
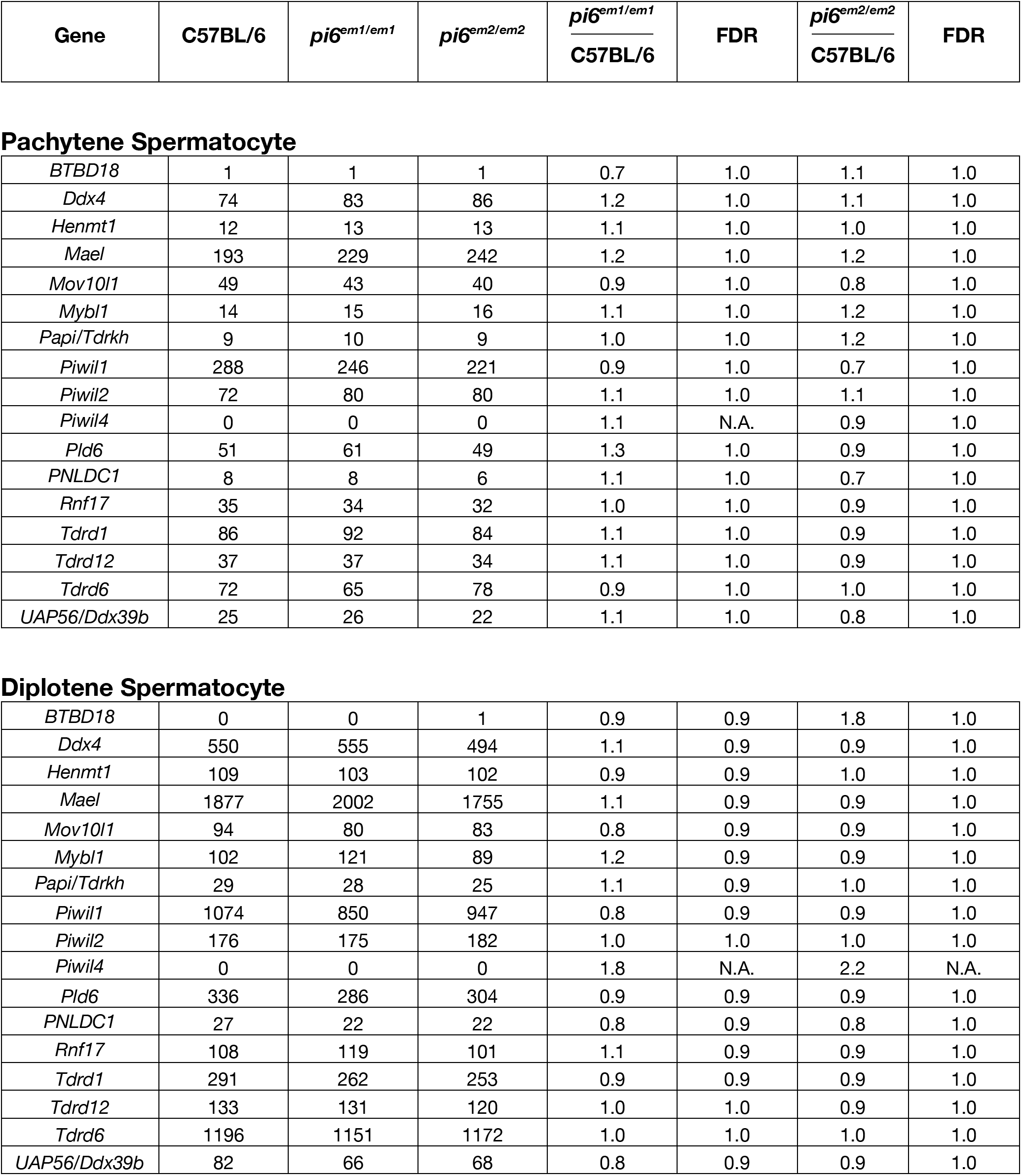
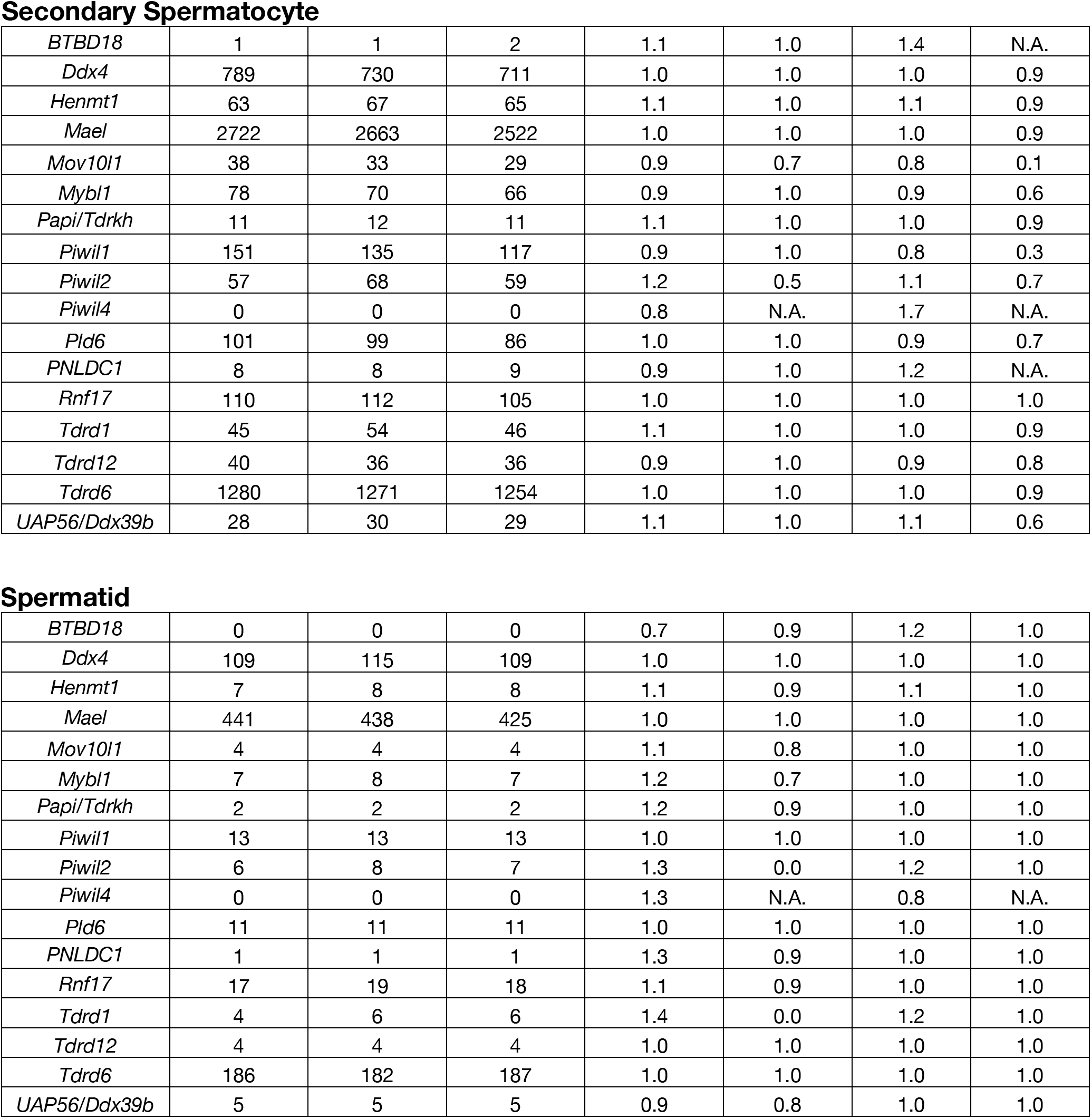
Abundance of piRNA pathway genes in *pi6^em1/em1^* and *pi6^em2/em2^* cells. Related to Figure 7 and S6. Mean abundance of piRNA genes (molecules per cell) in C57BL/6 versus *pi6^em1/em1^* and *pi6^em2/em2^* cells of replicate RNA-seq datasets. Changes in transcript abundance were calculated using DESeq2. Significant changes were required to increase or decrease >2-fold with FDR <0.05.

| Gene | C57BL/6 | $pi6^{em1/em1}$ | $pi6^{em2/em2}$ | $\frac{pi6^{em1/em1}}{C57BL/6}$ | FDR | $\frac{pi6^{em2/em2}}{C57BL/6}$ | FDR |
| --- | --- | --- | --- | --- | --- | --- | --- |
| Pachytene Spermatocyte |  |  |  |  |  |  |  |
| *BTBD18* | 1 | 1 | 1 | 0.7 | 1.0 | 1.1 | 1.0 |
| *Ddx4* | 74 | 83 | 86 | 1.2 | 1.0 | 1.1 | 1.0 |
| *Henmt1* | 12 | 13 | 13 | 1.1 | 1.0 | 1.0 | 1.0 |
| *Mael* | 193 | 229 | 242 | 1.2 | 1.0 | 1.2 | 1.0 |
| *Mov10l1* | 49 | 43 | 40 | 0.9 | 1.0 | 0.8 | 1.0 |
| *Mybl1* | 14 | 15 | 16 | 1.1 | 1.0 | 1.2 | 1.0 |
| *Papi/Tdrkh* | 9 | 10 | 9 | 1.0 | 1.0 | 1.2 | 1.0 |
| *Piwil1* | 288 | 246 | 221 | 0.9 | 1.0 | 0.7 | 1.0 |
| *Piwil2* | 72 | 80 | 80 | 1.1 | 1.0 | 1.1 | 1.0 |
| *Piwil4* | 0 | 0 | 0 | 1.1 | N.A. | 0.9 | 1.0 |
| *Pld6* | 51 | 61 | 49 | 1.3 | 1.0 | 0.9 | 1.0 |
| *PNLDC1* | 8 | 8 | 6 | 1.1 | 1.0 | 0.7 | 1.0 |
| *Rnf17* | 35 | 34 | 32 | 1.0 | 1.0 | 0.9 | 1.0 |
| *Tdrd1* | 86 | 92 | 84 | 1.1 | 1.0 | 0.9 | 1.0 |
| *Tdrd12* | 37 | 37 | 34 | 1.1 | 1.0 | 0.9 | 1.0 |
| *Tdrd6* | 72 | 65 | 78 | 0.9 | 1.0 | 1.0 | 1.0 |
| *UAP56/Ddx39b* | 25 | 26 | 22 | 1.1 | 1.0 | 0.8 | 1.0 |
| Diplotene Spermatocyte |  |  |  |  |  |  |  |
| *BTBD18* | 0 | 0 | 1 | 0.9 | 0.9 | 1.8 | 1.0 |
| *Ddx4* | 550 | 555 | 494 | 1.1 | 0.9 | 0.9 | 1.0 |
| *Henmt1* | 109 | 103 | 102 | 0.9 | 0.9 | 1.0 | 1.0 |
| *Mael* | 1877 | 2002 | 1755 | 1.1 | 0.9 | 0.9 | 1.0 |
| *Mov10l1* | 94 | 80 | 83 | 0.8 | 0.9 | 0.9 | 1.0 |
| *Mybl1* | 102 | 121 | 89 | 1.2 | 0.9 | 0.9 | 1.0 |
| *Papi/Tdrkh* | 29 | 28 | 25 | 1.1 | 0.9 | 1.0 | 1.0 |
| *Piwil1* | 1074 | 850 | 947 | 0.8 | 0.9 | 0.9 | 1.0 |
| *Piwil2* | 176 | 175 | 182 | 1.0 | 1.0 | 1.0 | 1.0 |
| *Piwil4* | 0 | 0 | 0 | 1.8 | N.A. | 2.2 | N.A. |
| *Pld6* | 336 | 286 | 304 | 0.9 | 0.9 | 0.9 | 1.0 |
| *PNLDC1* | 27 | 22 | 22 | 0.8 | 0.9 | 0.8 | 1.0 |
| *Rnf17* | 108 | 119 | 101 | 1.1 | 0.9 | 0.9 | 1.0 |
| *Tdrd1* | 291 | 262 | 253 | 0.9 | 0.9 | 0.9 | 1.0 |
| *Tdrd12* | 133 | 131 | 120 | 1.0 | 1.0 | 0.9 | 1.0 |
| *Tdrd6* | 1196 | 1151 | 1172 | 1.0 | 1.0 | 1.0 | 1.0 |
| *UAP56/Ddx39b* | 82 | 66 | 68 | 0.8 | 0.9 | 0.9 | 1.0 |
| Secondary Spermatocyte |  |  |  |  |  |  |  |
| *BTBD18* | 1 | 1 | 2 | 1.1 | 1.0 | 1.4 | N.A. |
| *Ddx4* | 789 | 730 | 711 | 1.0 | 1.0 | 1.0 | 0.9 |
| *Henmt1* | 63 | 67 | 65 | 1.1 | 1.0 | 1.1 | 0.9 |
| *Mael* | 2722 | 2663 | 2522 | 1.0 | 1.0 | 1.0 | 0.9 |
| *Mov10l1* | 38 | 33 | 29 | 0.9 | 0.7 | 0.8 | 0.1 |
| *Mybl1* | 78 | 70 | 66 | 0.9 | 1.0 | 0.9 | 0.6 |
| *Papi/Tdrkh* | 11 | 12 | 11 | 1.1 | 1.0 | 1.0 | 0.9 |
| *Piwil1* | 151 | 135 | 117 | 0.9 | 1.0 | 0.8 | 0.3 |
| *Piwil2* | 57 | 68 | 59 | 1.2 | 0.5 | 1.1 | 0.7 |
| *Piwil4* | 0 | 0 | 0 | 0.8 | N.A. | 1.7 | N.A. |
| *Pld6* | 101 | 99 | 86 | 1.0 | 1.0 | 0.9 | 0.7 |
| *PNLDC1* | 8 | 8 | 9 | 0.9 | 1.0 | 1.2 | N.A. |
| *Rnf17* | 110 | 112 | 105 | 1.0 | 1.0 | 1.0 | 1.0 |
| *Tdrd1* | 45 | 54 | 46 | 1.1 | 1.0 | 1.0 | 0.9 |
| *Tdrd12* | 40 | 36 | 36 | 0.9 | 1.0 | 0.9 | 0.8 |
| *Tdrd6* | 1280 | 1271 | 1254 | 1.0 | 1.0 | 1.0 | 0.9 |
| *UAP56/Ddx39b* | 28 | 30 | 29 | 1.1 | 1.0 | 1.1 | 0.6 |
| Spermatid |  |  |  |  |  |  |  |
| *BTBD18* | 0 | 0 | 0 | 0.7 | 0.9 | 1.2 | 1.0 |
| *Ddx4* | 109 | 115 | 109 | 1.0 | 1.0 | 1.0 | 1.0 |
| *Henmt1* | 7 | 8 | 8 | 1.1 | 0.9 | 1.1 | 1.0 |
| *Mael* | 441 | 438 | 425 | 1.0 | 1.0 | 1.0 | 1.0 |
| *Mov10l1* | 4 | 4 | 4 | 1.1 | 0.8 | 1.0 | 1.0 |
| *Mybl1* | 7 | 8 | 7 | 1.2 | 0.7 | 1.0 | 1.0 |
| *Papi/Tdrkh* | 2 | 2 | 2 | 1.2 | 0.9 | 1.0 | 1.0 |
| *Piwil1* | 13 | 13 | 13 | 1.0 | 1.0 | 1.0 | 1.0 |
| *Piwil2* | 6 | 8 | 7 | 1.3 | 0.0 | 1.2 | 1.0 |
| *Piwil4* | 0 | 0 | 0 | 1.3 | N.A. | 0.8 | N.A. |
| *Pld6* | 11 | 11 | 11 | 1.0 | 1.0 | 1.0 | 1.0 |
| *PNLDC1* | 1 | 1 | 1 | 1.3 | 0.9 | 1.0 | 1.0 |
| *Rnf17* | 17 | 19 | 18 | 1.1 | 0.9 | 1.0 | 1.0 |
| *Tdrd1* | 4 | 6 | 6 | 1.4 | 0.0 | 1.2 | 1.0 |
| *Tdrd12* | 4 | 4 | 4 | 1.0 | 1.0 | 1.0 | 1.0 |
| *Tdrd6* | 186 | 182 | 187 | 1.0 | 1.0 | 1.0 | 1.0 |
| *UAP56/Ddx39b* | 5 | 5 | 5 | 0.9 | 0.8 | 1.0 | 1.0 |

### Legends to Movies

**Movies S1-10. *pi6^em1/em1^* sperm motility.**

**Movie S1.** C57BL/6 sperm motility at 10 min.

**Movie S2.** *pi6^em1/em1^* sperm motility at 10 min.

**Movie S3.** C57BL/6 sperm motility at 90 min.

**Movie S4.** *pi6^em1/em1^* sperm motility at 90 min.

**Movie S5.** C57BL/6 sperm motility at 3 h.

**Movie S6.** *pi6^em1/em1^* sperm motility at 3 h

**Movie S7.** C57BL/6 sperm motility at 4 h.

**Movie S8.** *pi6^em1/em1^* sperm motility at 4 h.

**Movie S9.** C57BL/6 sperm motility at 5 h.

**Movie S10.** *pi6^em1/em1^* sperm motility at 5 h.

